# NAMPT activation uncovers a senescence-specific vulnerability and promotes healthy aging in combination with NAM

**DOI:** 10.64898/2026.08.06.743365

**Authors:** Michael Alcaraz, Raghuveer Ramachandra, Rouven Arnold, Marcos Garcia-Teneche, Adarsh Rajesh, Laurence Haddadin, Mey Taing, Xue Lei, Armin Gandhi, Theophilos Tzaridis, Karl Miller, Jessica Proulx, Amirhossein Nayeri Rad, Andrew Davis, Angela Liou, Hiroshi Tanaka, Tumpa Dutta, Rebecca Poritt, Valentin Cracan, Colin Loweth, Steven Olson, Stephen J. Gardell, Michael Jackson, Peter D. Adams

## Abstract

Aging is driven by multiple interacting processes, suggesting that effective strategies to promote healthy aging may require simultaneous targeting of more than one underlying mechanism. Here we identify a strategy that couples restoration of nicotinamide adenine dinucleotide (NAD+) homeostasis with selective targeting of senescent cells, two mechanistically linked features of aging. Senescent cells express elevated intracellular levels of nicotinamide phosphoribosyltransferase (NAMPT), the rate-limiting enzyme in the nicotinamide (NAM) salvage pathway for NAD+ biosynthesis. Despite increased NAMPT abundance, isotope-tracing studies revealed decreased NAD+ biosynthesis and consumption, indicating that elevated NAMPT abundance was not accompanied by a corresponding increase in NAD+ metabolic flux. Treatment with the NAMPT activator SBI-0802162 engaged the spare enzymatic capacity of NAMPT in senescent cells and produced a marked rise in intracellular NAD+ that, when sustained, disrupted their transcriptional program and selectively reduced the viability of senescent cells but not proliferating cells. In mice, SBI-0802162 reduced circulating NAM levels, suggesting that sustained NAMPT activation may be limited by substrate availability. This observation prompted the development of a combination approach using SBI-0802162 together with dietary NAM supplementation. Co-administration of SBI-0802162 and NAM robustly increased tissue NAD+, suppressed select age-associated inflammatory signatures and markers of cellular senescence in a tissue-specific manner. These molecular effects occurred alongside preserved physical performance in aged mice and reductions in food intake and body weight, which were observed whether SBI-0802162 was present in the chow or administered by oral gavage. Together, these findings establish a mechanistically integrated approach to target two convergent features of aging, NAD+ dysregulation and senescent cell accumulation, and support combined NAMPT activation and NAM supplementation as a strategy to promote healthy aging.

## INTRODUCTION

Aging is thought to be driven by the accumulation of multiple interacting forms of cellular damage and dysfunction, suggesting that effective interventions to promote healthy aging may need to address more than one process simultaneously^1^. Among these are altered nicotinamide adenine dinucleotide (NAD+) homeostasis and the accumulation of senescent cells, two mechanistically linked features of aging that may be effectively targeted in combination^2^.

Nicotinamide adenine dinucleotide (NAD+) is a central metabolic cofactor involved in redox reactions and a required co-substrate for NAD+-dependent enzymes including sirtuins (SIRTs), mono- and poly-ADP ribose polymerases (PARPs), ADP-ribose synthases (CD38/CD157), and sterile alpha toll/interleukin-1 receptor motif-containing 1 (SARM1)^3^. Through these enzymes, NAD+ influences energy metabolism, DNA repair, chromatin remodeling, immune regulation, cellular signaling, and senescence^4–6^. Mammalian cells maintain NAD+ through several biosynthetic routes. NAD+ can be synthesized de novo from tryptophan, from nicotinic acid (NA) through the Preiss-Handler pathway, or by the recycling of nicotinamide (NAM), a byproduct from NAD+-consuming enzymes, through the NAM salvage pathway. In most mammalian tissues, NAM salvage is the predominant source of NAD+^7–9^, with nicotinamide phosphoribosyltransferase (NAMPT) catalyzing the rate-limiting conversion of NAM and phosphoribosyl pyrophosphate (PRPP) to nicotinamide mononucleotide (NMN), which is then converted into NAD+ by one of three nicotinamide mononucleotide adenylyltransferases (NMNAT1-3)^7^.

NAD+ levels have been reported to decrease with age in multiple tissues^8, 10, 11^, and the decline has been linked to metabolic, neurodegenerative, and inflammatory disorders^12–15^. As a result, restoring NAD+ has become an attractive strategy to counter age-associated dysfunction. NAD+ precursor supplementation has shown benefits in preclinical models of multiple age-associated diseases^15–19^, but the extent to which these effects translate into durable functional improvements in humans remains uncertain. This may reflect limitations in precursor bioavailability, tissue-specific uptake, and uncertainty over which administered precursors best reach target tissues^20, 21^. Excessive intake of NAD+ precursors may also lead to accumulation of intermediates or byproducts that reshape tissue signaling and stress responses. For example, excessive NAD+ precursor supplementation can cause accumulation of N-methyl-2-pyridone-5-carboxamide (2PY) and N-methyl-4-pyridone-5-carboxamide (4PY), two metabolites associated with cardiovascular and renal pathologies^22–24^. These limitations highlight the need for alternative approaches that can more directly, safely, and durably restore or reprogram NAD+ metabolism *in vivo*.

The decline in NAD+ with age occurs alongside the accumulation of senescent cells, a cell state characterized by stable proliferative arrest that is associated with chronic inflammation and tissue dysfunction^25–34^. Although senescence plays beneficial roles in development, tumor suppression, and tissue repair, senescent cell accumulation with age can disrupt tissue structure and function through the chronic secretion of inflammatory mediators known as the senescence-associated secretory phenotype (SASP)^35–41^. Selective targeting of senescent cells can improve age-associated phenotypes in preclinical models, but the heterogeneity and context-dependence of senescence suggest that additional senotherapeutic strategies will be needed to fully realize the potential of this approach. Senescence is closely linked to NAD+ metabolism^25–34^. NAMPT and NAD+ have been implicated in SASP regulation through the AMPK-p53-p38-NF-κB axis^42^, and NAMPT itself has been reported to be upregulated and secreted as part of the SASP^43^. Senescent cells can also shape the tissue environment in ways that promote NAD+ depletion by recruiting and polarizing macrophages toward a CD38-high inflammatory state^44^.

Since altered NAD+ metabolism and cellular senescence are two interconnected regulators of aging, we set out to explore this relationship in more detail. Here, we show that in addition to upregulating and secreting NAMPT, senescent cells retain a substantial intracellular NAMPT pool. Despite this increase in NAMPT abundance, tracer studies revealed reduced incorporation of labeled NAM into NAD+, indicating diminished NAMPT-mediated NAD+ biosynthesis and suggesting a disconnect between NAMPT protein abundance and enzymatic output. To probe this disconnect, we utilized SBI-0802162, an orally bioavailable NAMPT activator and optimized analog of SBI-797812 with improved pharmacokinetic and pharmacodynamic properties^45–47^. By activating NAMPT, SBI-0802162 supports enhanced NAD+ biosynthesis through the NAM salvage pathway.

In senescent cells, pharmacologic activation of NAMPT with SBI-0802162 engaged the spare NAMPT enzymatic capacity and drove a marked increase in intracellular NAD+ that, when sustained, disrupted the normal stress-associated transcriptional program and selectively reduced senescent cell viability. Consistent with enhanced NAM utilization by the salvage pathway, *in vivo* dosing of SBI-0802162 reduced circulating NAM levels, suggesting that sustained NAMPT activation by SBI-0802162 may become limited by substrate availability. Dietary NAM supplementation together with SBI-0802162 mitigated the reduction in circulating NAM, enhanced tissue NAD+ elevation, and suppressed age-associated inflammatory and senescence-associated signatures in aged mice. These molecular effects were accompanied by attenuated age-associated frailty progression and preserved motor performance in aged mice, as well as reduced food intake and decreased body weight. The decreases in food intake and body weight were neither due to voluntary avoidance of compound-containing chow nor associated with other signs of malaise. Together, these findings identify decreased NAD+ turnover as a targetable metabolic vulnerability in senescent cells and support combining NAMPT activation with dietary NAM as a strategy to restore NAD+ homeostasis and counter age-associated inflammation.

## RESULTS

### Accumulation of NAMPT in senescent cells is accompanied by reduced rates of NAD+ synthesis and consumption

To define the changes in NAMPT expression during senescence, RNA and protein were collected from IMR90 primary human fibroblasts over a 21-day period following senescence induction by ionizing radiation (IR). Cellular senescence was confirmed by downregulation of CCNA2, CCNB1, lamin-B1 and phospho-pRb, as well as increased expression of CCND1, CCND2, CDKN2A/p16Ink4a, CDKN1A and several SASP genes (IL1α, IL1β, and IL8) as shown by immunoblot and/or qPCR^48^ (**Figure 1A** and **S1A**). Immunoblot and qPCR analysis also showed that NAMPT expression increased by day 12 post-IR and remained elevated through day 21 (**Figure 1A** and **S1A**). NAMPT upregulation was not limited to IR-induced senescence, as increased NAMPT expression was also observed in oncogene-induced senescence (OIS; **Figure 1B** and **S1B**), as well as in cells rendered senescent by etoposide or doxorubicin treatment (**Figure S1C**). Analysis of publicly available RNA-sequencing datasets further showed that NAMPT upregulation occurs in several senescence models, although the magnitude of this response varied across cell types and senescence-inducing stimuli (**Figure S1D**). These results are consistent with prior studies describing NAMPT as part of the SASP^42, 43^. However, immunofluorescence analysis showed that senescent cells also retain NAMPT intracellularly (**Figure 1C**), suggesting that increased NAMPT expression may have cell-intrinsic consequences in addition to its secreted role.

**Figure 1:**
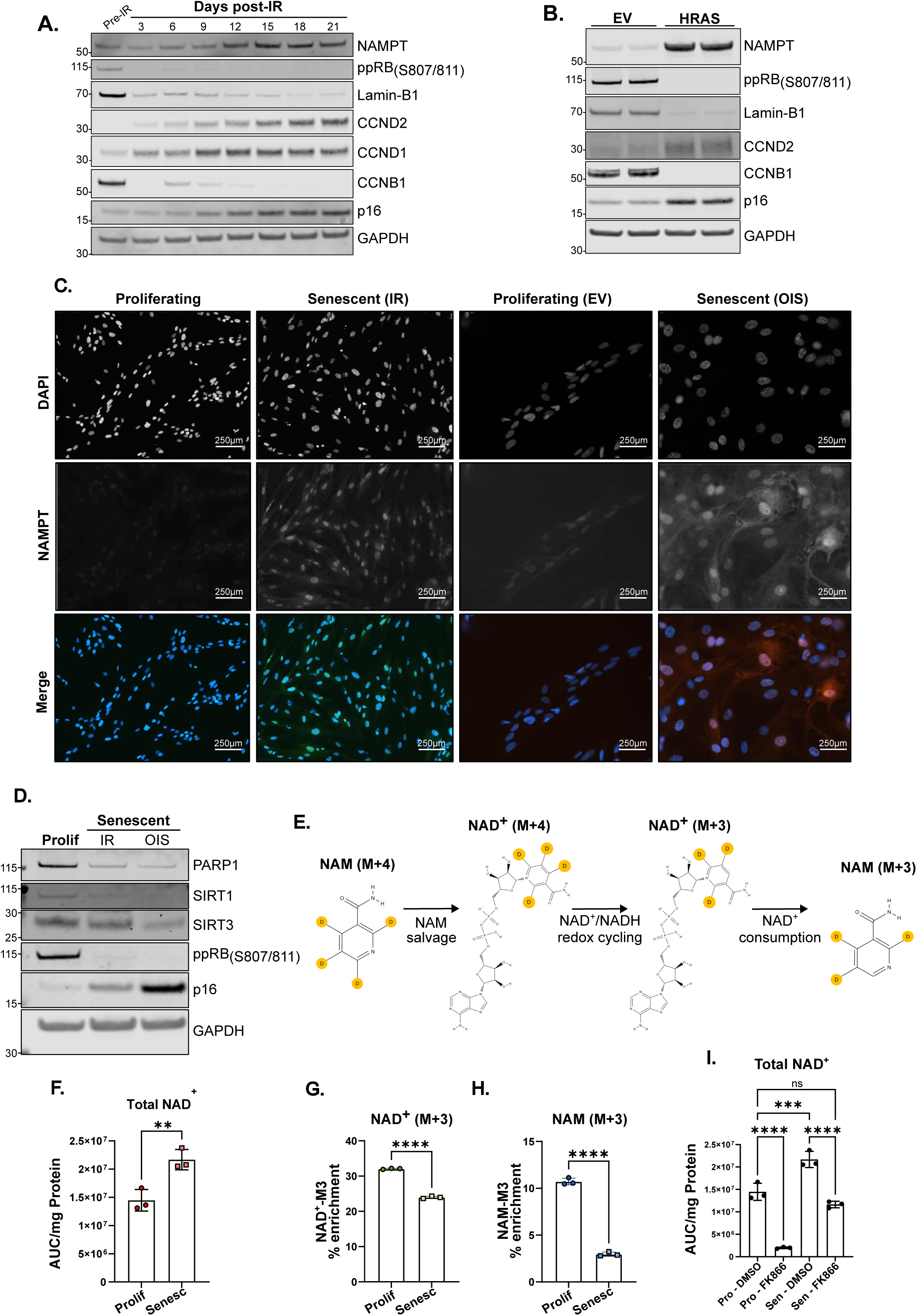
Accumulation of NAMPT in senescent cells is accompanied by reduced rates of NAD+ synthesis and consumption. **A**, Immunoblot of IMR90 cells prior to ionizing radiation and at the indicated timepoints post-irradiation. **B**, Immunoblot of IMR90 cells induced to undergo senescence by transduction with HRAS^G12V^ lentivirus (OIS). Protein was isolated 10 days post-transduction. **C**, Representative immunofluorescence images of proliferating, IR-induced senescent, proliferating transduced with empty vector (EV), and OIS IMR90 cells stained with DAPI or an NAMPT-targeting antibody. **D.** Immunoblot of major NAD+-consuming enzymes in proliferating, irradiation-induced and oncogene-induced senescent IMR90 cells. **E**, Diagram depicting the metabolic fate of [2,4,5,6-d_4_] NAM (NAM M+4) following cellular uptake. **F**, Total NAD+, NAD+ M+3 (**G**), and NAM M+3 (**H**) levels in proliferating and senescent (IR) IMR90 cells after exposure to NAM M+4 for 18 hours. **I**, Total NAD+ in proliferating and senescent cells after treatment with the NAMPT inhibitor FK866 (0.1 µM) for 18 hours. Each dot represents a biological replicate. Error bars denote mean ± s.d. Statistical analysis for 1F-1H was performed using unpaired parametric T-tests. Statistical analysis for 1I was performed using One-Way ANOVA and Dunnett’s multiple comparisons test. ns = not significant, *p<0.05, **p<0.01, ***p<0.001, ****p<0.0001.

To determine whether increased NAMPT abundance reflected a broader rewiring of NAD+ metabolism in senescent cells, we examined expression levels of major NAD+-consuming enzymes. Although RNA-level changes in these genes were roughly evenly divided between up and downregulated expression, protein levels of PARP1, SIRT1, and SIRT3, three major NAD+ consumers involved in DNA repair, cellular stress responses, and mitochondrial function, were reduced in both IR- and oncogene-induced senescent cells (**Figure 1D** and **S1E**). The reduced abundance of these consumers suggested a potential disconnect between elevated NAMPT levels and the apparent demand for NAD+ regeneration in senescent cells. We therefore directly assessed NAD+ biosynthetic and consumptive fluxes using stable isotope tracing by adding [2,4,5,6-d4] NAM (NAM M+4) to the media of IMR90 cells. The metabolic fate of NAM M+4 following cellular uptake is outlined in **Figure 1E**. NAM M+4 enters the salvage pathway and generates NAD+ M+4. Following synthesis, NAD+/NADH redox cycling results in loss of the deuterium at the C4 position of the nicotinamide ring, generating NAD+ M+3. Thus, the accumulation of NAD+ M+3 provides a readout of newly synthesized NAD+. Subsequent cleavage of the labeled NAD+ by NAD+-consuming enzymes releases NAM M+3, whose appearance provides a readout of consumption of the newly synthesized labeled NAD+ pool. Consistent with prior studies^42,43^, senescent cells exhibited a modest elevation of steady-state NAD+ levels despite the large increase in NAMPT (**Figure 1F**). Despite the increase in the steady-state NAD+ abundance, isotopic tracing revealed a marked reduction in the rates of both NAD+ biosynthesis and consumption in senescent cells. NAD+ biosynthesis was reduced as shown by a lower level of NAD+ M+3 within the total NAD+ pool, while NAD+ consumption was diminished as reflected by decreased abundance of NAM M+3 within the total NAM pool (**Figure 1G** and **1H**). Decreased NAD+ consumption in senescent cells was further confirmed by their attenuated response to NAMPT inhibition. Treatment of proliferating IMR90 cells with the NAMPT inhibitor FK866 (0.1 µM for 18 hours) depleted intracellular NAD+ by more than 90%, consistent with rapid NAD+ consumption driven by NAD+ consumers. In contrast, the NAD+ levels in senescent cells subjected to the same FK866 treatment declined by only approximately 50%, indicating slower NAD+ utilization (**Figure 1I**).

Together, these findings identify a unique metabolic state in which senescent cells exhibit elevated intracellular NAMPT and steady-state total NAD+ levels, yet paradoxically display decreased NAD+ biosynthesis and consumption. This suggests a disconnect between NAMPT abundance and enzymatic output that may leave spare catalytic capacity available for engagement.

### Sustained NAMPT activation disrupts mitochondrial and transcriptional homeostasis in senescent cells and selectively reduces their viability

The findings in **Figure 1** suggested that, although NAD+ biosynthetic flux is modestly reduced in senescent cells, elevated intracellular NAMPT may provide latent catalytic capacity that remains available for pharmacologic activation. Given the interest in strategies to elevate NAD+ as potential therapeutic interventions, we tested whether this capacity could be engaged by SBI-0802162, also known as 12c or DS68702229^45^, a small molecule NAMPT activator derived from SBI-797812^46^ (**Figure 2A**). We compared its effects with those produced by nicotinamide riboside (NR), an NAMPT-independent NAD+ precursor, in proliferating, irradiation-induced, and oncogene-induced senescent cells. NR treatment (500 µM, 24 hours) modestly increased intracellular NAD+ levels in both proliferating and senescent cells with 1.9- and 1.2-fold increases, respectively. In contrast, SBI-0802162 treatment (10 µM, 24 hours) produced a larger response in senescent cells, increasing NAD+ by 2.7-fold compared with a 1.9-fold increase in proliferating cells (**Figure 2B**). This differential response was also reflected in the NAD+/NADH ratio, which was significantly higher in senescent cells after SBI-0802162 treatment than after NR treatment (**Figure 2C**), consistent with an amplified response to NAMPT activation in senescent cells with elevated NAMPT abundance. Because NAD+ and NADH are central to mitochondrial redox balance and bioenergetic metabolism, we next asked whether the amplified NAD+ response to SBI-0802162 in senescent cells was accompanied by changes in mitochondrial homeostasis. To test this, we assessed mitochondrial content and membrane potential in proliferating and OIS cells following a 96-hour treatment with SBI-0802162, the NAMPT inhibitor FK866 (0.1 µM), or NR. In proliferating cells, neither SBI-0802162 nor NR had any significant effect on mitochondrial area or TMRM-based measures of mitochondrial membrane potential (either total intensity or intensity per mitochondrial area), whereas FK866 reduced TMRM intensity, as expected following NAMPT inhibition^49, 50^ (**Figure S2A**). In contrast, OIS cells showed a distinct response to NAMPT modulation. SBI-0802162 increased mitochondrial area per cell, but this increase was associated with a strong trend toward a reduction of TMRM intensity per unit area of the mitochondrial compartment, suggesting a dissociation between mitochondrial content and function (**Figure 2D**). Consistent with these results, Seahorse analysis revealed a reduction in both basal and maximal mitochondrial respiration elicited by SBI-0802162, as reflected by lower oxygen consumption rates (OCR; **Figure 2E** and **2F**). In contrast, extracellular acidification rate (ECAR) was largely unchanged under the same treatment conditions (**Figure 2G**). Treatment with SBI-0802162 did not affect the OCR or ECAR of proliferating IMR90 cells (**Figure S2B**), indicating that NAMPT activation suppressed mitochondrial respiratory activity selectively in senescent cells.

**Figure 2:**
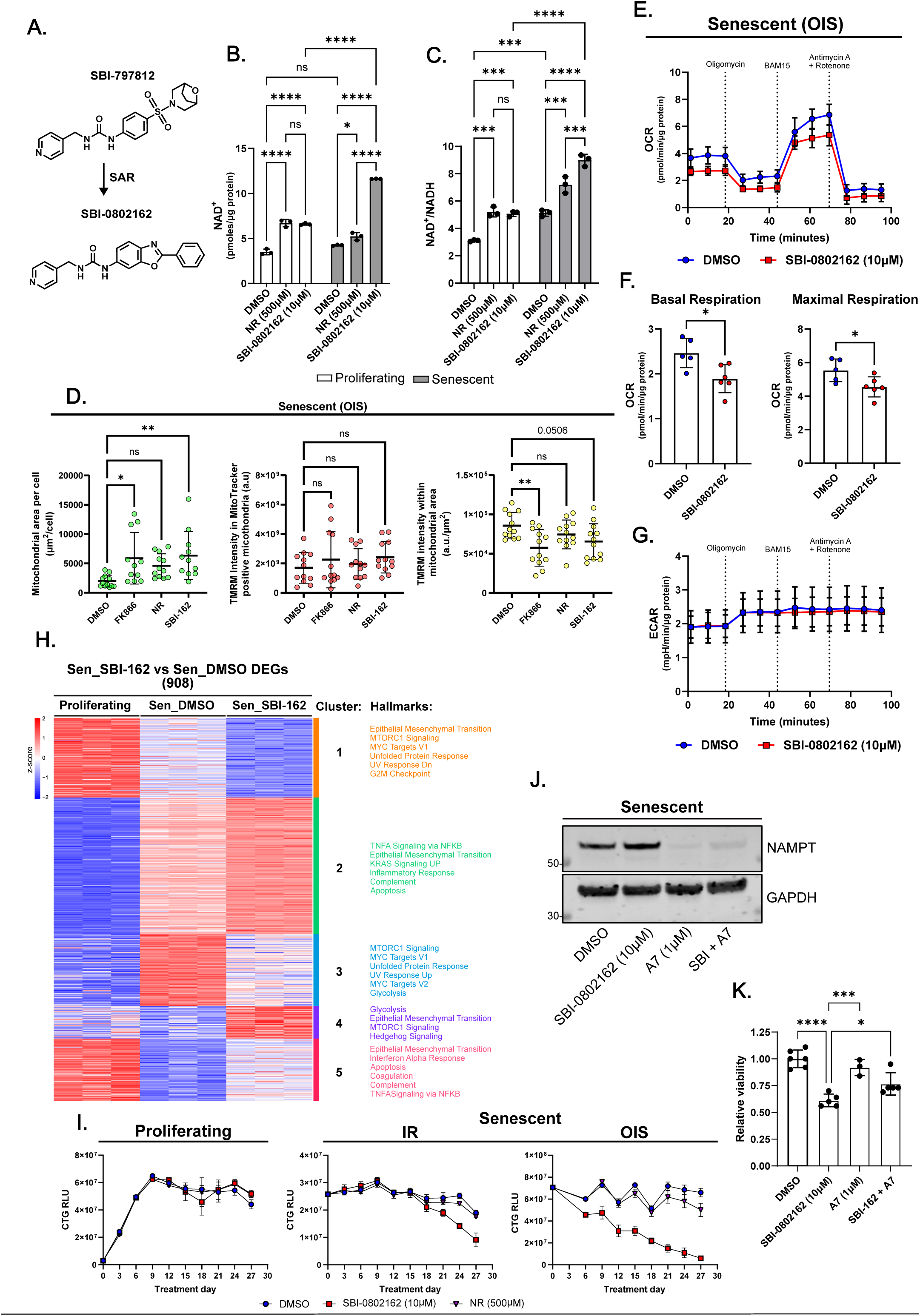
Sustained NAMPT activation disrupts mitochondrial and transcriptional homeostasis in senescent cells and selectively reduces their viability. **A**, Chemical structures of SBI-797812 and SBI-0802162, two NAMPT activators. **B**, NAD^+^ and NAD+/NADH ratio (**C**) in proliferating and irradiation-induced senescent IMR90 cells after treatment with nicotinamide riboside (NR; 500 µM) or SBI-0802162 (10 µM) for 24 hours. Metabolite levels were determined by enzymatic cycling assay. **D**, Mitochondrial area and membrane potential analysis in oncogene-induced senescent IMR90 cells after treatment with FK866 (0.1 µM), NR (500 µM), or SBI-0802162 (10 µM) for 96 hours. **E**, Seahorse mitochondrial stress test OCR trace in oncogene-induced senescent IMR90 cells treated with DMSO or SBI-0802162 (10 µM) for 96 hours. Results are normalized to total protein. **F**, Quantification of basal (left) and maximal (right) respiration from OCR measurements in 2E. Results are normalized to total protein. **G**, ECAR trace from Seahorse mitochondrial stress assay. Results are normalized to total protein. **H**, Heatmap of differentially expressed genes between SBI-0802162 and DMSO-treated oncogene-induced senescent cells, shown alongside proliferating cells for comparison. Major enriched Hallmark pathways within each gene cluster are indicated to the left of the heatmap. **I**, Longitudinal CellTiter-Glo (CTG) viability measurements in proliferating, irradiation-induced, and oncogene-induced senescent cells. X-axis shows CTG relative luminescent units (RLU). **J**, NAMPT protein levels in senescent cells treated with DMSO, SBI-0802162 (10 µM), the NAMPT PROTAC A7 (1 µM), and combined SBI-0802162 and A7. **K**, Relative viability of senescent cells treated with DMSO, SBI-0802162 (10 µM), A7 (1 µM), or combined SBI-0802162 and A7. Each dot in plots represents a biological replicate, except in Figure 2D, where each dot represents a single frame; 3 biological replicates were analyzed. Error bars denote mean ± s.d. Statistical analysis was performed using One-Way ANOVA and Dunnett’s multiple comparisons test. ns = not significant, *p<0.05, **p<0.01, ***p<0.001, ****p<0.0001.

We next asked whether these mitochondrial changes were accompanied by broader changes in the senescent state. Given the role of NAD+ as a link between metabolic and transcriptional programs^51^, we sought to determine the transcriptomic consequences in OIS cells after 96 hours of SBI-0802162 treatment. Consistent with the expected transcriptional shift during senescence, proliferating and OIS cells showed broad transcriptional changes by PCA, with 11,346 differentially expressed genes (DEGs; FDR < 0.05, **Figure S2C** and **S2D, left**). As expected, senescent cells showed increased expression of genes related to SASP and p53 signaling, including NF-κB and inflammatory response programs, as well as decreased expression of genes associated with cell cycle, including E2F targets, G2/M checkpoint, and mitotic spindle genes (**Figure S2E**). By comparison, 908 DEGs were identified between DMSO- and SBI-0802162-treated OIS cells. Of these genes, 420 reflected an exacerbation of the normal senescence response, including 288 that were upregulated and 132 downregulated by SBI-0802162 (**Figure S2D, right**). The 288 genes upregulated in senescence and further activated by SBI-0802162 were associated with TNFα signaling via NF-κB, inflammatory response, coagulation, complement, and IL6-JAK-STAT3 signaling (**Figure 2H**, cluster 2 and **S2F**), while the 132 genes downregulated in senescence and further suppressed by SBI-0802162 were associated with MYC targets, unfolded protein response, mTORC1 signaling, and G2/M checkpoint (**Figure 2H** cluster 1, and **S2F**). In addition, 372 genes were regulated by SBI-0802162 in a manner antagonistic to the normal senescence program. This included 233 genes that were increased in OIS cells and reduced by SBI-0802162 (**Figure 2H**, cluster 3) and 139 genes that were decreased in OIS and increased by SBI-0802162 treatment (**Figure 2H**, cluster 5). These oppositely regulated genes were associated with mTORC1 signaling, MYC targets, unfolded protein response, glycolysis, and EMT. The remaining 116 DEGs represented a treatment-specific response and were induced by SBI-0802162. These genes were associated with glycolysis, EMT, mTORC1, and Hedgehog signaling (**Figure 2H**, cluster 4). Together, these findings indicate that SBI-0802162 neither clearly reinforced nor suppressed the senescence-associated transcriptional program but instead dysregulated it by amplifying some features of senescence, opposing others, and inducing an additional treatment-specific program. Notably, this response was not restricted to OIS, as IR-induced senescent cells exposed to the same treatment regimen exhibited many of the same transcriptional changes observed in OIS cells (**Figure S2G**).

Given the seemingly detrimental mitochondrial and transcriptional dysregulation observed after SBI-0802162 treatment in senescent cells, we next asked whether sustained NAMPT activation selectively compromised senescent cell viability. To test this, we treated proliferating, irradiation-induced, and oncogene-induced senescent cells with DMSO, SBI-0802162, or NR and monitored viability over 27 days with media and compound replenishment every third day. NR was included as a comparator because it increases NAD+ without directly engaging the elevated NAMPT pool in senescent cells (**Figure 2B**, **2C**). Proliferating cells maintained viability after treatment with either SBI-0802162 or NR. In contrast, SBI-0802162 reduced the viability of both IR-induced and OIS cells, whereas NR had no effect on the viability of either senescent model (**Figure 2I**). This preferential effect on senescent cells was further supported by cell counts, which showed a time-dependent decline in senescent cell number after SBI-0802162 treatment, while proliferating cell number remained largely unchanged compared with DMSO-treated controls (**Figure S2H**). To determine whether the loss of viability was mediated by NAMPT rather than off-target activity of SBI-0802162, we used the PROTAC A7^52^ to deplete NAMPT in IR-induced senescent cells. We assessed viability following treatment with A7 (1 µM) or SBI-0802162 alone, or after NAMPT depletion by A7 before and during SBI-0802162 exposure. SBI-0802162 alone significantly reduced senescent cell viability, whereas A7 alone had no effect. However, depletion of NAMPT with A7 attenuated the reduction in viability caused by SBI-0802162, indicating that elevated NAMPT is required for the full senolytic effect of SBI-0802162 (**Figure 2J** and **2K**).

Collectively, these results show that elevated NAMPT abundance combined with decreased NAD+ consumption sensitizes senescent cells to NAMPT activation, resulting in an amplified NAD+ response. Sustained NAD+ elevation in senescent cells by SBI-0802162 disrupts mitochondrial homeostasis, dysregulates the senescence-associated transcriptional program, and selectively compromises senescent cell viability, consistent with senolytic activity.

### NAMPT activation combined with NAM supplementation prevents substrate limitation and more effectively elevates NAD+ in mouse tissues

Having established the NAD+-boosting effects and reduced senescent-cell viability after SBI-0802162 treatment *in vitro*, we next evaluated its impact on NAD+ levels *in vivo*. To define the *in vivo* pharmacodynamic profile of SBI-0802162, 3-month-old male C57BL/6J fed a control diet (containing nicotinic acid at 30 mg kg^-1^ of chow) were treated by oral gavage with a single dose of 30 or 100 mg kg^-1^ SBI-0802162, and plasma was collected 1.5 hours later for analysis. These doses were selected based on prior studies showing NAD+-boosting activity across mouse tissues with no adverse effects^45,46^. SBI-0802162 treatment significantly reduced plasma NAM levels, an effect consistent with activation of the enzymatic function of NAMPT (**Figure 3A**). However, this observation suggested that sustained NAMPT activation may create a substrate-limited state that could hinder the efficacy of SBI-0802162 *in vivo* and motivated the development of a combination strategy using SBI-0802162 and dietary NAM supplementation. To test if dietary NAM supplementation could mitigate the plasma NAM depletion caused by SBI-0802162, 3-month-old male C57BL/6J mice were fed a control (lacking NAM but containing nicotinic acid) or NAM-supplemented diet (1 g kg^-1^ chow) for two days before receiving vehicle or SBI-0802162 at 30 or 100 mg kg^-1^ by oral gavage, with plasma collected 1.5 hours later (**Figure 3B**). The NAM dose was selected based on a prior study^53^. LC-MS analysis showed that NAM supplementation increased circulating NAM by approximately 10-fold relative to vehicle-treated mice on control chow. Although SBI-0802162 induced a dose-dependent reduction in plasma NAM, NAM levels in supplemented animals never fell below those of vehicle-treated mice on control chow, even at the 100 mg kg^-1^ dose (**Figure 3C**), indicating that dietary NAM supplementation preserves plasma NAM after treatment with high doses of SBI-0802162.

**Figure 3.**
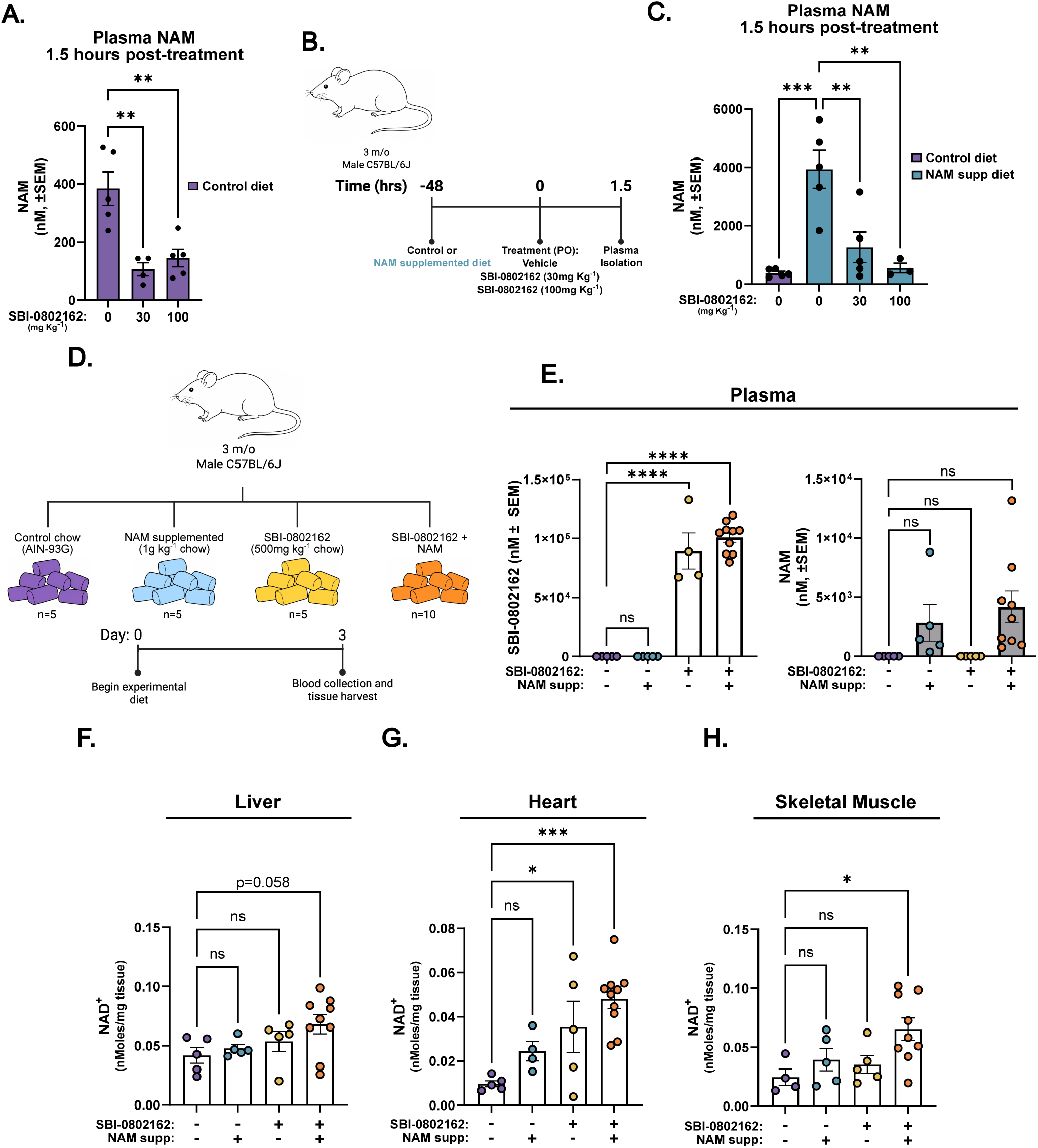
NAMPT activation combined with NAM supplementation prevents substrate limitation and more effectively elevates NAD+ in mouse tissues. **A**, Plasma NAM levels measured 1.5 hours after a single oral dose of vehicle or SBI-0802162 at 30 or 100 mg kg^-1^ in mice maintained on a control (AIN-93G) diet. **B**, Experimental design for acute pharmacodynamic assessment of SBI-0802162 with or without NAM supplementation. **C**, Plasma NAM levels 1.5 hours after treatment in mice maintained on NAM-supplemented diet and treated with vehicle or SBI-0802162 at 30 or 100 mg kg^-1^. **D**, Experimental design to assess the effect of in-chow NAM, SBI-0802162, or combined SBI-0802162 and NAM on NAD+ levels. **E**, Plasma levels of SBI-0802162 and NAM after 3 days of dietary treatment. **F**, Tissue NAD+ levels in liver, heart (**G**), and skeletal muscle (**H**) following 3 days of dietary treatment with control chow or diets containing NAM, SBI-0802162, or a combination of both agents. Each dot in plots represents an individual mouse. Error bars represent mean ± SEM. Statistical analysis was performed using One-Way ANOVA and Dunnett’s multiple comparisons test. ns = not significant, *p<0.05, **p<0.01, ***p<0.001, ****p<0.0001.

To determine whether NAM supplementation could enhance the tissue NAD+ response to SBI-0802162, healthy 3-month-old male C57BL/6J mice were fed custom diets containing SBI-0802162 (500 mg kg^-1^ chow, corresponding to approximately 100 mg kg^-1^ body weight per day), NAM (1 g kg^-1^ chow), or both agents for 3 days, after which plasma, liver, skeletal muscle, and heart were collected for analysis in the early morning (**Figure 3D**). All diets were well tolerated, with no significant changes in body weight or overt signs of toxicity over the 3 days (**Figure S3A**). LC-MS/MS confirmed abundant systemic exposure of SBI-0802162 in treated groups, which was not impacted by NAM supplementation (**Figure 3E, left**). Circulating NAM levels also trended higher in NAM-supplemented groups, although the increase did not reach statistical significance (**Figure 3E, right**). Tissue NAM levels were largely unchanged in the liver and increased in the heart and skeletal muscle following combination treatment (**Figure S3B-S3D**), an effect consistent with the rapid absorption, utilization, and excretion of NAM by tissues^54, 55^. Despite the limited changes in tissue NAM, the combination of SBI-0802162 with dietary NAM produced an upward trend in liver NAD^+^, whereas SBI-0802162 or NAM alone had little effect (**Figure 3F**). Remarkably, in skeletal muscle and heart, the combination increased NAD+ levels above those in controls and those achieved with either agent alone (**Figure 3G** and **3H**).

### Aging induces shared inflammatory transcriptional programs across tissues that are differentially reversed by NAM, SBI-0802162, or their combination

Given the established role of NAD+ in gene regulation^3^, its reported age-associated decline^8, 10, 11^, prior studies showing benefits of NAD+ supplementation^5^, and the *in vitro* senolytic effect reported here, we next set out to test whether NAM, SBI-0802162, or their combination could modify age-associated transcriptomic changes. To address this, 20-month-old C57BL/6J mice were fed diets containing NAM, SBI-0802162, or both agents for 21 days, after which liver, skeletal muscle, and heart were collected for further analysis (**Figure 4A**). These tissues were selected based on the NAD+-boosting effects observed in the short-term dietary study in **Figure 3** and because they represent major sites of age-associated metabolic and functional decline that could benefit from NAD+ restoration. Mass spectrometry analysis of plasma confirmed systemic exposure to SBI-0802162 and NAM in mice fed their respective diets (**Figure S4A**).

**Figure 4.**
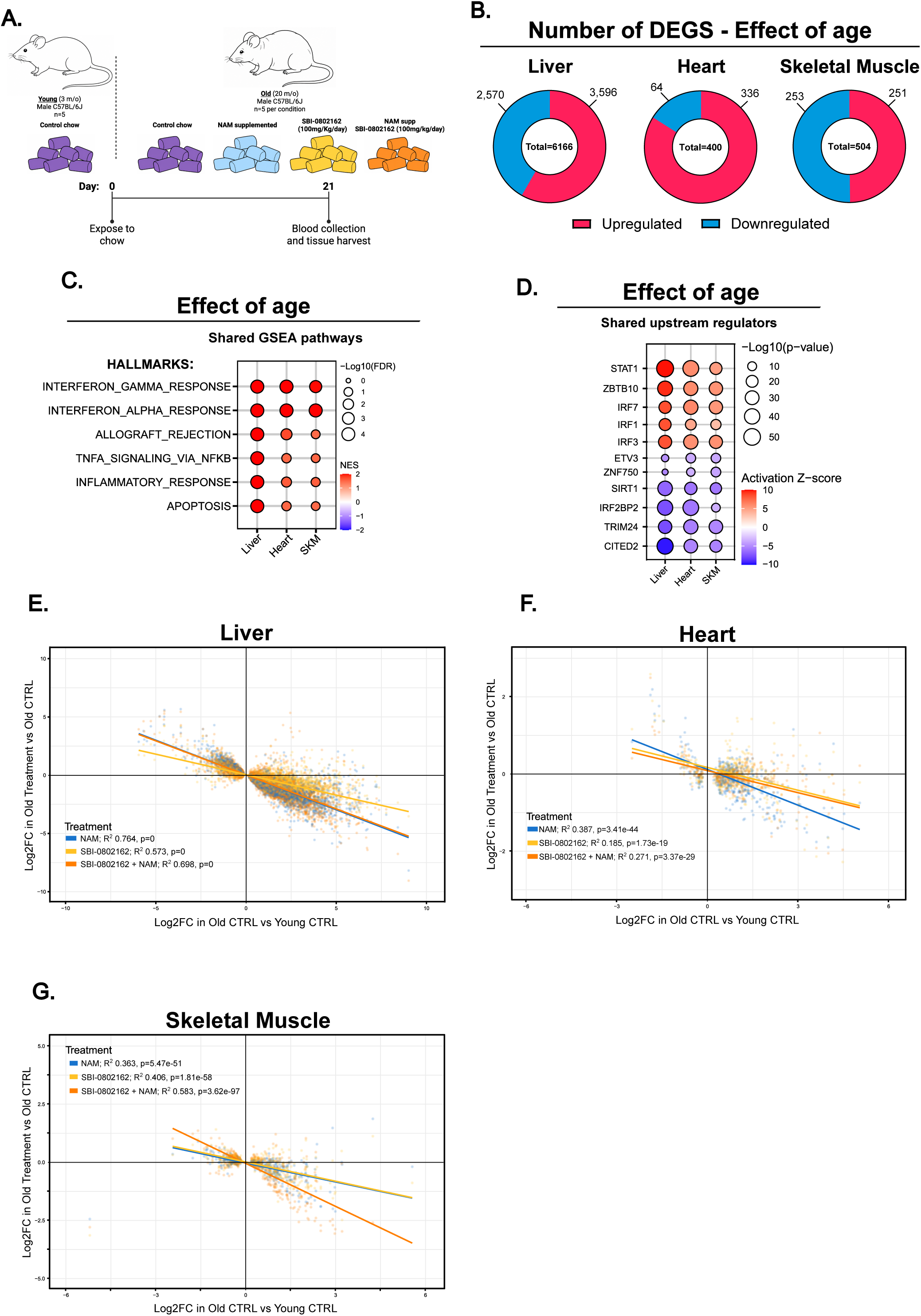
Aging induces shared inflammatory transcriptional programs across tissues that are differentially reversed by NAM, SBI-0802162, or their combination. **A**, Experimental design to address transcriptomic effects after 21 days of in-chow treatment with NAM, SBI-0802162, and combined SBI-0802162 plus NAM in aged mice. **B**, Donut plots showing the number and direction of age-associated differentially expressed genes (DEGs; FDR < 0.05) in liver, heart, and skeletal muscle. **C**, Shared GSEA hallmark pathways enriched with age across liver, heart, and skeletal muscle. **D**, Shared upstream regulators affected with age across liver, heart and skeletal muscle. **E**, Correlation analysis comparing age-associated transcriptional changes with treatment-induced changes of the same genes in liver, heart (**F**), and skeletal muscle (**G**). Each dot in plots represents an individual mouse. Error bars represent mean ± SEM. Statistical analysis was performed using One-Way ANOVA and Dunnett’s multiple comparisons test. ns = not significant, *p<0.05, **p<0.01, ***p<0.001, ****p<0.0001.

Unbiased transcriptomic analysis revealed clear tissue-specific differences in the extent of age-associated gene expression changes. Consistent with PCA plots showing the clearest separation between young and aged liver samples, liver had the largest number of age-associated DEGs (FDR < 0.05) with 6,166 genes significantly altered with age (3,596 upregulated and 2,570 downregulated; **Figure 4B, left** and **S4B**). Heart and skeletal muscle showed more modest age-associated transcriptional changes, with 400 (336 upregulated and 64 downregulated) and 504 (251 upregulated and 253 downregulated) DEGs, respectively (**Figure 4B** middle, **4B,** right, **S4C**, and **S4D**). Despite these quantitative differences, GSEA identified inflammatory pathways, particularly interferon (IFN) response programs, as shared features of aging across all three tissues (**Figure 4C**), as observed previously^56–62, 87^. Consistent with this, Ingenuity Pathway Analysis (IPA) predicted activation of STAT1 and multiple IFN regulatory factors (IRFs) with age, whereas SIRT1, a NAD+ deacetylase linked to metabolic regulation, healthspan, and longevity^63–65, 87^, was among the top transcriptional regulators predicted to be inhibited in aged tissues (**Figure 4D**).

We next examined the effects of NAM, SBI-0802162, or their combination on the aging transcriptome. For each tissue we focused on genes significantly altered with age (FDR < 0.05) and compared the direction and magnitude of the age-associated change with the treatment-associated change for the same genes. This approach allowed us to determine whether each intervention tended to oppose or reinforce the transcriptional changes that occurred with age. Across liver, heart, and skeletal muscle, treatment-associated gene expression generally opposed age-associated transcriptional changes, as reflected by negative regression slopes in all three tissues (**Figure 4E-4G**). However, the strength of this effect varied by tissue and intervention. In the liver, NAM-containing interventions, either alone or in combination with SBI-0802162, produced stronger age-opposing effects than SBI-0802162 alone (**Figure 4E**). In the heart, the negative slopes also indicated partial opposition of age-associated changes, although NAM produced a stronger age-opposing effect than SBI-0802162 alone or the combination. Of note, the regression lines were offset from the origin, suggesting that the treatment response was not simply a proportional reversal of aging and may include additional heart-specific transcriptional effects (**Figure 4F**). Strikingly, in skeletal muscle, combined SBI-0802162 and NAM produced the clearest age-opposing effect, exceeding either intervention alone (**Figure 4G**).

Altogether, these findings are consistent with previous reports^56–62, 87^ showing shared upregulation of inflammatory and IFN-response signatures across aged tissues and indicate that NAD+-enhancing interventions can shift aged tissues toward a more youthful transcriptional state, although the extent of this restoration is highly tissue-and intervention-dependent. Specifically, NAM alone and in combination with SBI-0802162 were most effective in liver, NAM alone in heart, and the combination was the most effective intervention in skeletal muscle.

### NAMPT activation combined with dietary NAM supplementation suppresses age-associated inflammatory and IFN-response programs in a tissue-dependent manner

After assessing the overall extent to which each intervention opposed age-associated transcriptional changes, we next examined the specific genes and pathways altered by each intervention across the aged liver, heart, and skeletal muscle. Marked tissue-specific responses were observed, with the combination of NAM and SBI-0802162 producing its most pronounced and coordinated transcriptional effects in skeletal muscle.

In the aged liver, NAM alone produced the largest transcriptional response, significantly altering 4,220 genes, with 3,414 opposing their age-associated change. SBI-0802162 alone produced a smaller response, altering 964 genes, including 769 that opposed their age-associated expression change. Combined SBI-0802162 and NAM produced an intermediate response, significantly altering 3,420 genes, with 2,507 shifting in the opposite direction of their age-associated change (FDR < 0.05; **Figure 5A**). In all three cases, only very few age-associated changes were enhanced by the interventions (7, 4, or 10 genes, as indicated in **Figure 5A**). Pathway analysis revealed that NAM alone and in combination with SBI-0802162 broadly suppressed age-associated inflammatory and IFN-response programs, whereas SBI-0802162 alone produced a more modest effect (**Figure 5B**). Consistent with this, IPA showed that both NAM-containing interventions reversed the predicted activation status of upstream regulators associated with inflammation, including STAT1 and multiple IFN regulatory factors, while predicting activation of SIRT1, an effect consistent with other NAD+-boosting interventions^66, 87^. SBI-0802162 alone had no predicted effect on upstream regulators affected by age in the liver (**Figure 5C**). Targeted analysis of IFN-stimulated genes further supported these findings with both NAM-containing interventions suppressing ISG expression (**Figure 5D**). Consistent with the transcriptomic data, immunoblot analysis of liver lysates confirmed reduced total STAT1 and phospho-STAT1 (Y701) after NAM alone or in combination with SBI-0802162 (**Figure 5E**), indicating that suppression of the IFN-response program by NAM-containing interventions was also apparent at the protein level. Of note, by all measures, combined SBI-0802162 and NAM showed a trend toward greater suppression of age-associated IFN signaling and ISGs when compared to either agent alone (**Figure 5B-5E**).

**Figure 5.**
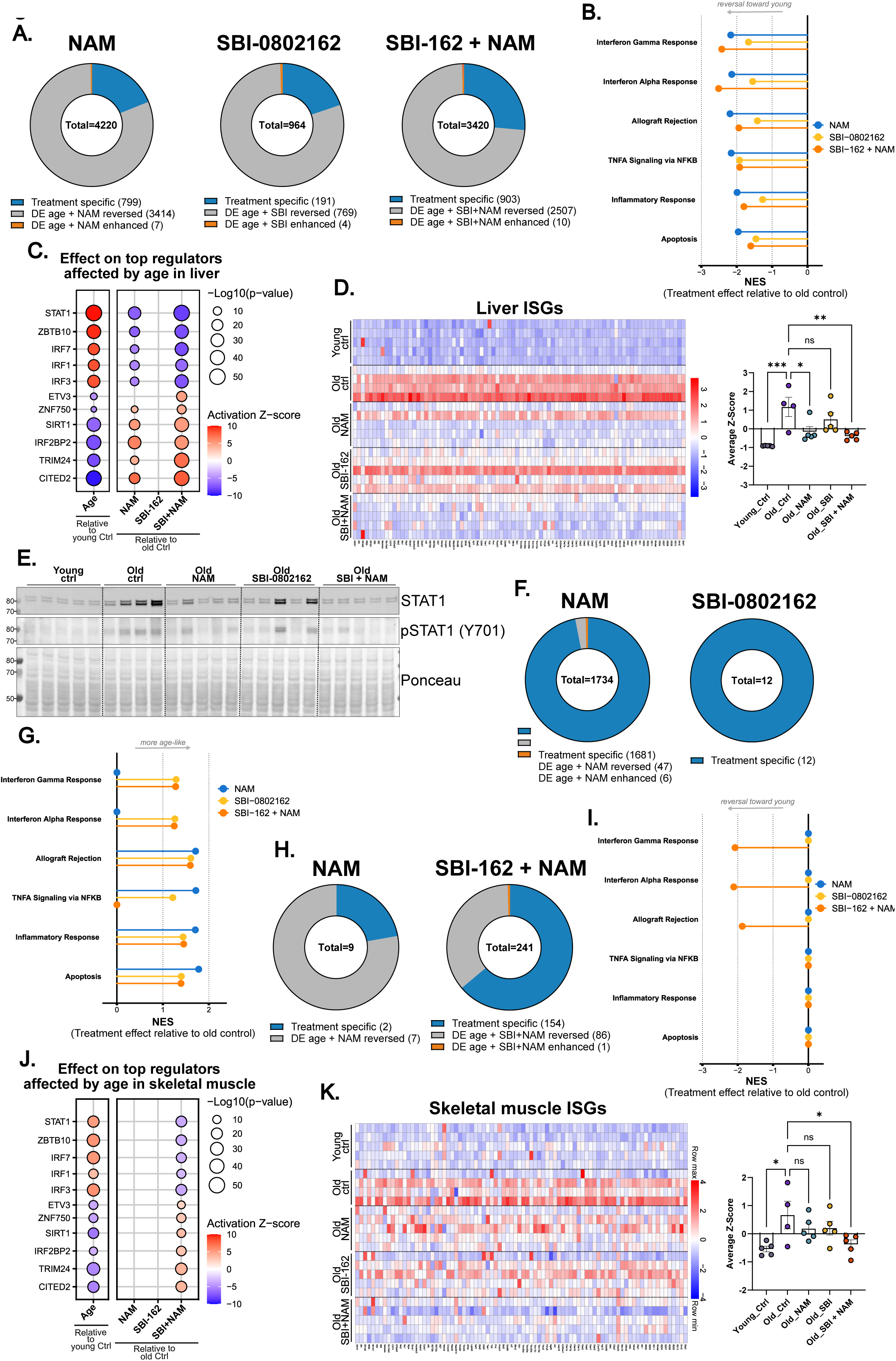
NAMPT activation combined with dietary NAM supplementation suppresses age-associated inflammatory and IFN-response programs in a tissue-dependent manner. **A**, Donut plots showing the number and classification of differentially expressed genes (DEGs; FDR < 0.05) relative to old controls in the aged liver following in-chow treatment with NAM, SBI-0802162, or combined SBI-0802162 plus NAM for 21 days. DEGs are categorized as treatment-specific, age-associated DEG and reversed by treatment, and age-associated DEG and enhanced with treatment. **B**, GSEA pathway analysis showing how NAM, SBI-0802162, and combined SBI-0802162 plus NAM modulate, in liver, commonly altered pathways by age across tissues of interest. **C**, Upstream regulator analysis showing how NAM, SBI-0802162, and combined SBI-0802162 plus NAM modulate, in liver, regulators commonly altered by age across tissues of interest. **D**, Heatmap of interferon-stimulated gene (ISG) expression in the liver. Quantification of average ISG Z-scores is to the right. **E**, Immunoblot analysis of STAT1 and phosphorylated STAT1 (Y701) in liver lysates from young and old control samples as well as treated aged mice. Each lane represents an individual mouse. **F**, Donut plots showing the number and classification of DEGs relative to old controls in the aged heart. **G**, Modulation of age-associated GSEA pathways in heart, focusing on pathways commonly altered by age across the tissues of interest. **H**, Donut plot showing the number and classification of DEGs relative to old controls in the aged skeletal muscle. **I**. GSEA pathway analysis showing how NAM, SBI-0802162, and combined SBI-0802162 plus NAM modulate, in skeletal muscle, commonly altered pathways by age across tissues of interest. **J**. Upstream regulator analysis showing how NAM, SBI-0802162, and combined SBI-0802162 plus NAM modulate, in skeletal muscle, upstream regulators commonly altered by age across tissues of interest. **K**, Heatmap of ISG expression in skeletal muscle. Quantification of average ISG Z-score is to the right. In plots 5D right, and 5K right each dot represents an individual mouse. Error bars represent mean ± SEM. Statistical analysis was performed using One-Way ANOVA and Dunnett’s multiple comparisons test. ns = not significant, *p<0.05, **p<0.01, ***p<0.001, ****p<0.0001.

The aged heart showed a qualitatively distinct and more complex response to the interventions. NAM alone altered 1,734 genes relative to aged controls, with 1,681 of these being treatment-specific and only 47 opposing their age-associated expression pattern (**Figure 5F**, **left**), indicating that the cardiac response to NAM largely reflected a treatment-specific transcriptional program rather than reversal of age-associated gene expression. The response of the aged heart to NAM included both upregulation and downregulation of metabolic genes (**Figure S5A, S5B**). By comparison, SBI-0802162 alone produced minimal transcriptomic changes, altering only 12 genes, none of which were affected by age (**Figure 5F**, **right**). The combination of SBI-0802162 and NAM produced no significant DEGs at the same threshold (FDR < 0.05). In contrast to the liver, and despite the modest reversal of age-associated transcriptomic changes observed in **Figure 4F**, pathway analysis showed positive enrichment of inflammatory pathways in all three treatment groups relative to age-matched controls (**Figure 5G**). Notably, the NAM-induced inflammatory and metabolic changes were attenuated in mice receiving combined SBI-0802162 and NAM (**Figure 5G**, **S5A**, and **S5B**).

Compared to the liver and heart, the aged skeletal muscle showed a more selective response to the NAD+-enhancing interventions. NAM alone produced 9 DEGs relative to aged controls, 7 of which shifted in the direction opposite to aging (**Figure 5H**, **left**). SBI-0802162 alone did not produce any significant DEGs at the same threshold (FDR < 0.05). In contrast, the combination of SBI-0802162 and NAM altered 241 genes, including 86 that opposed their age-associated expression changes and only one that enhanced them (**Figure 5H**, **right**). Pathway analysis showed that combined SBI-0802162 and NAM preferentially suppressed inflammatory and IFN-response programs in aged skeletal muscle, while changes induced by NAM or SBI-0802162 alone were not associated with any significant pathway enrichment (**Figure 5I**). IPA showed that the combination reduced the predicted activation of pro-inflammatory upstream regulators affected by age, including STAT1 and multiple IFN regulatory factors, while predicting activation of SIRT1 (**Figure 5J**). Targeted ISG analysis further supported this effect, showing significant suppression of ISG expression exclusively in mice receiving combined SBI-0802162 and NAM (**Figure 5K**).

Collectively, these findings highlight the tissue-specific nature of NAD+-enhancing interventions in aging. Although liver, heart, and skeletal muscle share an age-associated inflammatory signature, their responses to treatment were distinct. NAM alone suppressed inflammatory signaling in the liver, but had limited benefit in skeletal muscle, and heightened inflammatory transcriptional programs in the heart. By comparison, combined SBI-0802162 and NAM produced a more balanced cross-tissue transcriptional response, enhancing the anti-inflammatory effects of NAM in the aged liver, attenuating the inflammatory transcriptional remodeling induced by NAM in the heart, and driving the strongest anti-inflammatory response in the aged skeletal muscle.

### Combined NAMPT activation and dietary NAM supplementation reduce senescence-associated transcriptional programs and p16Ink4a protein levels in aged tissues

Because age-associated ISG expression has been linked to senescent cell accumulation^67^, and sustained NAMPT activation led to a senolytic effect *in vitro*, we asked whether the suppression of inflammatory programs observed after treatment with NAM, SBI-0802162, or their combination was accompanied by a decrease in senescence-associated markers in aged tissues.

To assess this, we first calculated average Z-scores across multiple senescence-related gene lists, including the SenDiego (curated by SenNet San Diego TMC), SenMayo, Fridman_Up, Casella_up, and CellAge signatures^68–71^. Aging significantly increased senescence-associated gene signature scores in liver, as expected. These scores were reduced most strongly by NAM, modestly by SBI-0802162, and to an intermediate extent by the combination of SBI-0802162 and NAM (**Figure 6A**). In contrast, the heart showed limited age-associated induction of these signatures and treatment with NAM or SBI-0802162 alone increased the senescence-associated scores relative to age-matched controls. Combined SBI-0802162 and NAM also increased these scores, but to a lesser extent than either single intervention (**Figure 6B**). In skeletal muscle, combined SBI-0802162 and NAM was the only intervention that robustly reduced the age-related increase in senescence-associated signature scores (**Figure 6C**), suggesting that treatment effects on senescence-associated transcriptional programs are highly tissue dependent. The senescence-suppressive effects of NAM, and particularly the combination of SBI-0802162 and NAM, were also reflected at the protein level by reduced p16Ink4a abundance in the liver. A similar reduction was observed in the kidney, another tissue with a high senescent cell burden with age as assessed by immunoblotting^72^ (**Figure 6D**, **6E**).

**Figure 6.**
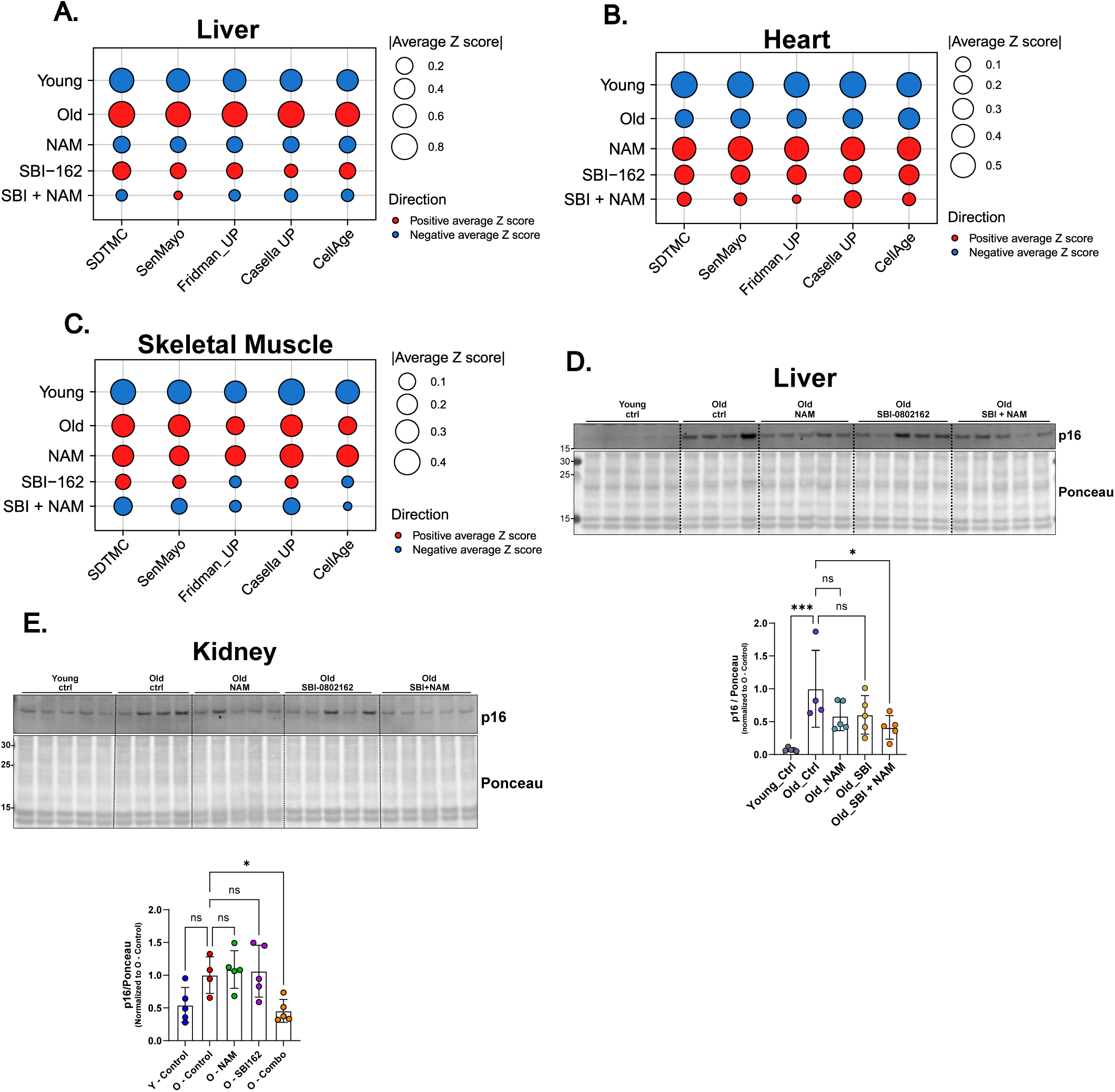
Combined NAMPT activation and dietary NAM supplementation reduce senescence-associated transcriptional programs and p16Ink4a protein levels in aged tissues. **A**, Dot plot showing average Z-scores for multiple senescence-associated gene signatures, including SDTMC, SenMayo, Fridman_UP, Casella_UP, and CellAge, in liver, heart (**B**), and sketal muscle (**C**) from young control, aged control, and aged mice treated with NAM, SBI-0802162, or combined SBI-0802162 plus NAM. Bubble size represents the magnitude of the average Z-score, and color indicates the direction of the signature score. **D**, immunoblot analysis and quantification of p16Ink4a protein levels in liver and kidney (**E**) from young control, aged control, and aged mice treated with the indicated interventions.

Together, these results show that the effects of NAM, SBI-0802162, and their combination on senescence-associated signatures broadly paralleled their effects on the wider aged transcriptome. NAM produced the strongest reduction of senescence-associated signatures in the liver. The combination of SBI-0802162 and NAM reduced these signatures in both liver and skeletal muscle, while partially blunting the senescence-associated transcriptional response observed with either single agent in the aged heart. Skeletal muscle showed the clearest and most consistent response to the combination of SBI-0802162 and NAM, with evidence of transcriptome rejuvenation, reduced inflammatory signaling, and suppression of senescence-associated transcriptional programs.

### SBI-0802162 combined with dietary NAM attenuates frailty progression while reducing body weight and food intake in mice

Because combined SBI-0802162 and NAM produced the most consistent NAD+-boosting effects across tissues (**Figure 3** and **S3**) and showed comparable or greater activity than the single agents on several key transcriptomic readouts, particularly in liver and skeletal muscle (**Figures 4-6**, **S4**, and **S5**), we selected the combination for a long-term functional study to determine whether these molecular effects translated into functional benefits during aging. To address this, naturally aged 18-month-old C57BL/6J mice were maintained on control diet or chow containing combined SBI-0802162 and NAM for 12 weeks, with sex-matched 3-month-old mice included as young controls. Frailty and motor coordination were assessed every 4 weeks using the mouse frailty index^73, 74^ and rotarod testing, respectively, while body weight and food consumption were monitored weekly (**Figure 7A**). Young and aged control-fed mice showed a progressive increase in frailty over the course of the study. In contrast, frailty scores remained stable in aged mice receiving the combination diet, consistent with attenuation of age-associated frailty progression (**Figure 7B** and **S6A**).

**Figure 7.**
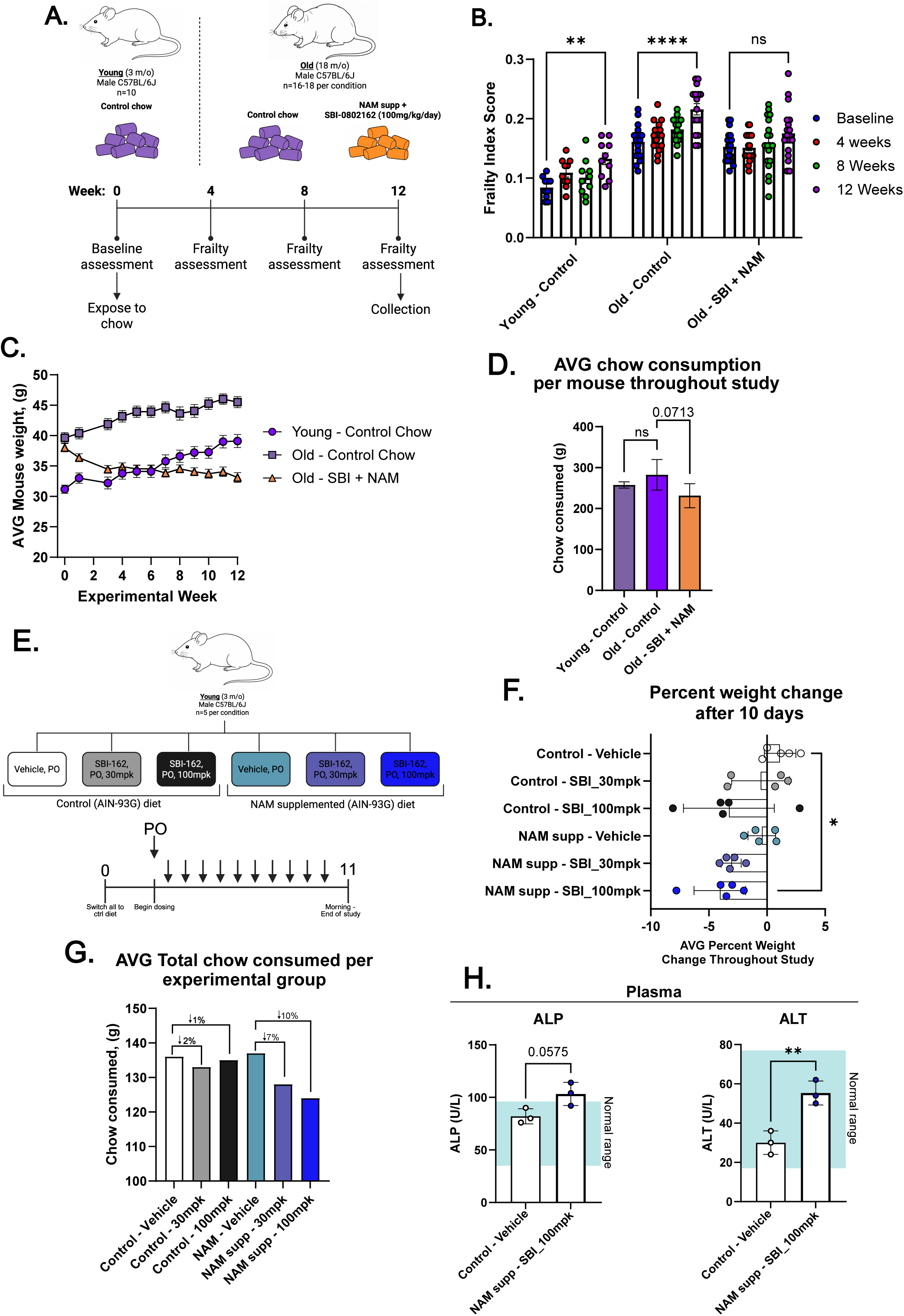
SBI-0802162 combined with dietary NAM attenuates frailty progression while reducing body weight and food intake in mice. **A**, Experimental design to address long-term functional consequences of dietary intervention with combined SBI-0802162 and NAM in aged mice. **B**, Longitudinal frailty index scores measured at baseline and after 4, 8, and 12 weeks of dietary intervention. Higher values indicate increased frailty. **C**, Average mouse body weight over the 12-week study. **D**, Average chow consumed per mouse over the 12-week study for each experimental condition (n=5 per experimental condition). **E**, Experimental design to address if delivery of SBI-0802162 independently of diet led to similar weight and food intake effects. **F**, Percent body weight change following 10 days of treatment in mice receiving the indicated diet and SBI-0802162 dosing regimens via oral gavage. **G**, Average total chow consumed per experimental group during the 10-day study (average consumption of 5 mice). Percent differences in food consumption between treatment groups are indicated. **H**, Plasma alkaline phosphatase (ALP) and alanine aminotransferase (ALT) levels following 10-day study. In plots 7B, 7F, and 7H each dot represents an individual mouse. Error bars represent mean ± SEM. Statistical analysis for 7H was performed using unpaired parametric T-tests. Statistical analysis for 7B was performed using two-way ANOVA with Tukey’s multiple comparisons test. Statistical analysis for 7F was performed using One-Way ANOVA and Dunnett’s multiple comparisons test. ns = not significant, *p<0.05, **p<0.01, ***p<0.001, ****p<0.0001.

Analysis of individual frailty components revealed that reduced body weight contributed substantially to the lower frailty scores observed in mice treated with combined SBI-0802162 and NAM (**Figure S6B**). Young and aged mice maintained on control diet gained weight over the course of the study, whereas aged mice receiving SBI-0802162 and NAM lost an average of 14% of their body weight during the first three weeks of treatment, after which body weight stabilized (**Figure 7C**). Of note, the reduction in body weight of mice on the combination diet was accompanied by a nonsignificant trend toward decreased food consumption (**Figure 7D**).

We first considered the possibility that reduced food intake could reflect decreased palatability of the compound-containing chow leading to voluntary dietary restriction. To investigate this, we asked whether decreased chow consumption was also observed when SBI-0802162 was delivered independently of the diet. Three-month-old male C57BL/6J mice were maintained on a control or a NAM-supplemented diet (1 g kg^-1^ chow) and treated daily with vehicle or SBI-0802162 at 30 or 100 mg kg^-1^ by oral gavage for 10 days (**Figure 7E**). Under these conditions, SBI-0802162 produced a nonsignificant, dose-dependent trend toward reduced body weight that became statistically significant when combined with dietary NAM supplementation (**Figure 7F**). NAM or SBI-0802162 alone had no effect on food intake, whereas combined SBI-0802162 and NAM caused a dose-dependent reduction in chow consumption (**Figure 7G**). These findings indicate that reduced food intake is not simply explained by decreased palatability of SBI-0802162-containing chow but instead reflects a pharmacologic effect of combined SBI-0802162 and NAM treatment.

Reduced food consumption could also reflect sickness or malaise induced specifically by the combination treatment. However, several observations argue against this interpretation. First, although plasma alanine aminotransferase (ALT) and alkaline phosphatase (ALP) were modestly increased in mice receiving 100 mg kg^-1^ SBI-0802162 plus NAM, both remained within the normal reference range, arguing against overt hepatotoxicity (**Figure 7H**). Second, mice treated with combined SBI-0802162 and NAM lacked other signs of malaise including tremors, gait abnormalities, or gastrointestinal issues (**Figure S6B**). Third, the lower frailty score observed in combination-fed mice was not explained by body weight alone, as other frailty components, including alopecia and piloerection, also improved (**Figure S6B**). Fourth, rotarod performance declined over time in aged control-fed mice as expected, but was preserved in aged mice receiving the combination diet, with a strong trend toward improved performance relative to aged controls (**Figure S6C**). Together, these observations do not support sickness or malaise as the cause of reduced food intake in the combination-fed mice.

Altogether, combined SBI-0802162 and dietary NAM supplementation attenuated age-associated frailty progression and preserved motor performance in aged mice. These benefits were accompanied by reduced food intake and decreased body weight, but occurred without evidence of voluntary avoidance of compound-containing chow, overt sickness, or loss of physical function.

## DISCUSSION

Here, we show that altered NAD+ metabolism creates a targetable vulnerability in senescent cells, linking two mechanistically related features of aging: disrupted NAD+ homeostasis and senescent cell accumulation. Our study identifies a distinct state of NAD+ metabolism in senescent cells in which NAD+ levels and NAMPT abundance are elevated, but NAD+ turnover is reduced. By engaging the excess enzymatic capacity of NAMPT, SBI-0802162 produced a large increase in NAD+ in senescent cells *in vitro* and selectively impaired their viability when treatment was sustained. *In vivo*, NAMPT activation depleted circulating NAM, motivating a combination strategy with dietary NAM supplementation to maintain substrate availability. Combined SBI-0802162 and NAM increased tissue NAD+ more effectively than either agent alone, reduced age-associated inflammatory and senescence-related programs in liver and skeletal muscle, attenuated frailty progression, and preserved physical performance in aged mice. Across both dietary and gavage-based dosing paradigms, combined SBI-0802162 and NAM also reduced food intake and body weight without evidence of overt toxicity or malaise.

The broader implications of these findings extend beyond the effects of SBI-0802162 and NAM on physiological aging. Strategies that preserve tissue function, suppress chronic inflammation, and limit the detrimental effects of senescent cells could have relevance not only for healthy aging but also for age-associated diseases characterized by impaired NAD+ metabolism and persistent inflammatory signaling, such as neurodegenerative and metabolic disorders as well as musculoskeletal diseases. Although NAD+ precursor supplementation has shown benefit in several preclinical settings, its efficacy is often limited by tissue-specific uptake, dependence on endogenous biosynthetic capacity, rapid precursor metabolism, and diversion into pathways that do not contribute directly to NAD+ production^75–78^. These limitations may render precursor supplementation insufficient in some tissues and may also produce unintended consequences through secondary metabolism. By combining NAM supplementation with activation of the rate-limiting enzyme in the salvage pathway, our approach simultaneously increases substrate availability and promotes its productive incorporation into NAD+.

An important unresolved question is why NAMPT remains elevated in senescent cells. Previous studies have shown that NAMPT expression and secretion increase during senescence^42,43^, and our findings extend those observations by showing that senescent cells also retain a substantial intracellular NAMPT pool. NAMPT induction could represent a program intended to maintain NAD+ biosynthetic capacity during persistent stress, DNA damage, or metabolic remodeling. In addition, NAMPT may be induced as part of the broader inflammatory and secretory program of senescence, with a portion of the protein subsequently released into the extracellular space. Despite elevated intracellular NAMPT levels, senescent cells showed reduced rates of NAD+ biosynthesis and consumption, together with lower levels of major NAD+-consuming enzymes including PARP1/2 and SIRT1/3. NAMPT is subject to feedback inhibition by NAD+, which was elevated in senescent cells. Accumulation of its product NMN can also inhibit forward NAMPT flux through product binding and reversal of the NAMPT reaction^47, 79^, raising the possibility that while intracellular NAMPT is abundant in senescence, it is functionally constrained by the accumulation of its pathway metabolites in this cellular state. SBI-0802162 and related activators may release this constraint by reducing feedback inhibition, thereby engaging the enzymatic capacity that is normally unavailable^46^.

A second unresolved question is how sustained NAMPT activation leads to loss of senescent cell viability. Sustained NAD+ elevation by SBI-0802162 dysregulated core features of the senescence program, including inflammatory signaling and cell-cycle-associated transcriptional programs. Because senescence is established in response to sublethal damage, further disruption of the pathways that maintain this state may impose additional stress and reduce the ability of senescent cells to remain viable. Because the K_m_ of several NAD+-consuming enzymes fall within the range of cellular NAD+ concentrations, their activity can remain sensitive to substrate availability. Thus, despite the reduced abundance of some PARPs, sirtuins, and other NAD+ consumers in senescent cells, the marked increase in NAD+ induced by SBI-0802162 could enhance the activity of the remaining enzyme pool and dysregulate NAD+-dependent signaling, chromatin regulation, and stress responses. One example is excessive PARP activity, which could increase energetic demand and promote pathways associated with cell death. Our findings also implicate mitochondrial dysfunction as a link between NAD+ overload, cellular stress, and impaired viability. We therefore propose that elevated NAMPT abundance, together with decreased NAD+ utilization, allows SBI-0802162 to drive NAD+ above a tolerable range in senescent cells. When sustained, this state disrupts metabolic homeostasis and eventually compromises viability. Determining whether mitochondrial dysfunction is the initiating event or a downstream consequence and identifying the form of cell death involved are important next steps.

The cardiac response further illustrates why NAD+-enhancing interventions must be evaluated in a tissue-specific manner. NAM alone induced a predominantly age-independent transcriptional program in the aged heart, which was attenuated when SBI-0802162 was co-administered. One possibility for these effects is that excess NAM that is not efficiently directed into NAM salvage is diverted by secondary metabolism into 1-methylnicotinamide (1-MNA), N-methyl-2-pyridone-5-carboxamide (2PY) and N-methyl-4-pyridone-5-carboxamide (4PY). Some of these metabolites have been associated with cardiovascular and renal pathology^22, 23^. Levels of these metabolites were not measured in this study, and this explanation remains speculative. Nevertheless, the cardiac findings suggest that increasing precursor availability without simultaneously increasing its productive use may have unintended consequences. In this context, NAMPT activation may not simply enhance the NAD+-boosting effect of NAM but may also redirect its metabolism and limit the potentially detrimental response produced by NAM supplementation alone.

Finally, the functional benefits of combined SBI-0802162 and NAM treatment occurred alongside an unexpected reduction in food intake and body weight. Weight loss plateaued three weeks into the study and remained stable thereafter. Treated mice showed no overt signs of illness, maintained physical performance, and exhibited a strong trend toward improved rotarod performance relative to aged controls. These observations argue against generalized toxicity as a cause of reduced food intake and weight loss. Moreover, administration of SBI-0802162 via oral gavage reproduced the reduction in body weight and chow intake, supporting a pharmacologic effect rather than reduced palatability of the treatment-containing diet. Although NAD+-enhancing interventions have been associated with reduced adiposity in preclinical models^45, 66, 80, 81^, the accompanying reduction in food intake suggests that NAMPT activation may also affect appetite or satiety. Determining whether this response originates from central appetite-regulating circuits, peripheral metabolic signals, or altered communication between tissues will require direct measurements of body composition, energy expenditure, feeding behavior, and satiety-related pathways.

Several limitations exist in our study. First, although prolonged NAMPT activation selectively reduced senescent cell viability *in vitro*, the exact mechanism underlying this effect remains unclear. *In vivo*, the reduction in p16Ink4a protein levels and senescence-associated transcriptional programs following combined SBI-0802162 and NAM treatment is consistent with a decrease in senescent cell burden^82, 86^, but does not directly demonstrate senescent cell elimination. Restricted sample availability prevented higher-resolution spatial and cell-type-specific analyses needed to distinguish senescent cell elimination from reduced expression of senescence-associated markers within tissue-resident cells. Second, our tissue NAD+ measurements are also limited by our reliance on steady-state metabolite measurements, which provide little information about NAD+ flux. The larger NAD+ increase observed in skeletal muscle and heart relative to liver may therefore reflect greater NAD+ accumulation in tissues with slower turnover, whereas rapid NAD+ utilization in the liver may buffer changes in steady-state abundance^75^. Measurements of NAD+ flux and secondary NAM metabolites would help clarify the basis of these tissue-specific responses and may also shed light on the distinct transcriptional effects observed in the aged heart. Third, combined SBI-0802162 and NAM was selected for the long-term studies in **Figure 7A-D** because it produced the most consistent NAD+ increases and showed comparable or greater activity than NAM alone on key transcriptomic readouts across tissues. However, the absence of single-agent groups in this study limits our ability to determine the contribution of each agent and whether the effects on frailty depended on the combination. Future studies comparing NAM, SBI-0802162, and the combination will therefore be needed. Finally, mouse body composition, energy expenditure, and satiety-related pathways were not directly assessed, limiting our ability to explain the reductions in food intake and body weight associated with treatment.

Collectively, our findings establish NAMPT activation as a strategy for exploiting a metabolic vulnerability in senescent cells and, when paired with NAM, a means for modifying multiple molecular and functional features of aging. Combined SBI-0802162 and NAM increased tissue NAD+, reduced inflammatory and senescence-associated programs in select aged tissues, preserved physical function in aged mice, and attenuated several of the transcriptional responses produced by NAM alone. Establishing the translational potential of this approach will require defining the mechanisms underlying its effects on senescent cells and inflammation, resolving the basis of the changes in food intake and body weight, and more rigorously assessing the consequences and safety of sustained NAD+ elevation across tissues.

## METHODS

### Cell culture and senescence induction

IMR90 primary human fibroblasts were obtained from the American Type Culture Collection (ATCC CCL-186) and grown at 37°C with 3.5% O_2_ and 5% CO_2_ in Dulbecco’s modified Eagle’s medium (DMEM; Gibco, 10313-121) supplemented with 10% FBS (Corning, 35-010-CV), 1% penicillin/streptomycin (Gibco, 15140-122) and 2mM glutamine (Gibco, 25030-081). Cells were continuously checked for mycoplasma contamination and were used at passages 12-30. Irradiation induced senescence was induced by 20-Gray x-ray irradiation of 1×10^6^ cells in a 10cm dish. Cells were split 3 days after irradiation and allowed to undergo the senescence process for 15 days unless otherwise noted. Media was replenished every third day. For etoposide-and doxorubicin-induced senescence, 80% confluent IMR90 cells were treated with 50mM etoposide or doxorubicin for 24 hours, after which compound was removed. Cells were split 3 days after treatment and allowed to undergo the senescence process for 10 days unless otherwise noted. For oncogene-induced senescence, IMR90 cells were transduced with lentiviral particles encoding HRAS G12V or GFP and selected with puromycin (1µg/mL) 24-hours post-transduction. Two-days post-transduction cells were passaged into a 7cm dish and were allowed to reach confluency for 3 days with puromycin selection. 5 days post-transduction, cells were passaged into final assay plasticware and cultured for an extra 5 days to allow for senescence establishment. Media was replenished every third day.

### Plasmids

pLenti CMV/TO RasV12 Puro and pLenti CMV GFP Puro (658-5) were obtained from Addgene (#22262 and #17448) and used for oncogene-induced senescent experiments.

### Lentivirus infection

Lentivirus was generated by transfecting 293T cells with expression vector, VSVG envelope vector, and psPAX2 packaging vector in lipofectamine 2000 (Invitrogen 52887). Virus was collected over 3 days and cleared for any cell debris by centrifugation. Virus was titrated to the minimum amount required for >90% viability after puromycin selection (1µg/mL for 5 days). Infections were performed overnight (16 hours), in the presence of 8µg/mL polybrene (Millipore TR-1003-G).

### In vitro treatments

Treatments with SBI-0802162 (Sundia chemicals), FK866 (Selleckchem S2799), nicotinamide riboside (NR; Selleckchem S2935), and A7 (Tocris Bioscience 7842/5) started on day 15-post irradiation for irradiation induced senescent cells or 10-days post-transduction for oncogene induced senescent cells unless otherwise noted. For long-term treatments media and compounds were replenished every third day. SBI-0802162 was used at 10 µM, FK866 was used at 0.1 µM, NR was used at 500 µM, and A7 was used at 1 µM.

### NAD^+^ tracer studies

Proliferating and senescent cells were maintained in Dulbecco’s modified Eagle’s medium (DMEM; Gibco, 10313-121) supplemented with 10% FBS (Corning, 35-010-CV), 1% penicillin/streptomycin (Gibco, 15140-122) and 2mM glutamine (Gibco, 25030-081) until start of tracing experiment and then switched for 18 hours to flux DMEM with treatment, if applicable. Briefly, custom Dulbecco’s modified Eagle’s medium lacking niacinamide and L-tryptophan was generated (Gibco) as was the base media for flux DMEM. Complete flux DMEM was generated by addition of 10% dialyzed FBS (ThermoFischer, 26400044), 2mM glutamine (Gibco, 25030-081), 1% penicillin/streptomycin (Gibco, 15140-122), L-tryptophan (to a final concentration of 78.4 µM, ThermoFischer, J62508.22), and a 50-50 mix of NAM (MilliporeSigma, N0636-100G) and NAM (D_4_) (Cambridge Isotope Labs, DLM-6883) to a final concentration of 32 µM. At start of experiment, cells were switched from DMEM to flux DMEM for 18 hours. After 18 hours, media was removed quickly and cells were washed with cold PBS (Gibco, 14190-144) containing 5 µM 78c (CD38 inhibitor; Selleckchem, S8960-5MG). PBS was quickly removed and liquid nitrogen was added to the dish. Liquid nitrogen was decanted and the frozen monolayer was scraped and transferred to a falcon tube. The complete harvesting procedure was performed on wet or dry ice. High resolution mass spectrometry was performed as previously described^83^.

### NAD^+^ enzymatic cycling assays

NAD^+^ enzymatic cycling assays were performed utilizing the NAD/NADH-Glo^TM^ (Promega, G9072) reagent according to the manufacturer’s protocol. Briefly, cells were washed with cold PBS (Gibco, 14190-144) and lysed in a 1% Dodecyltrimethylammonium bromide (DTAB, Sigma-Aldrich, D8638-25G) solution in NaOH. Lysate was then split into 2. Half of the lysate was treated with 0.4N HCl and heated to 60°C for 20 minutes. The second half was heated to 60°C for 20 minutes. Trizma® was then added to the half containing HCl, and a mixture of HCl and Trizma® was added to the second half of the lysate. Prepared NAD/NADH-Glo^TM^ reagent was added to both lysates and luminescent readings were obtained at 10, 20, 30, and 40 minutes post-addition of the luminescent reagent. NAD^+^ and NADH standards (Sigma, N8285 and N6660) were generated in the same solution as the experimental samples. Luminescence was recorded using a PHERAstar FSX plate reader (BMG Labtech). Concentrations of NAD^+^ and NADH was normalized to total protein quantified by Bradford assay (Thermo 23246).

### Cell viability assays

Cell viability was assessed utilizing the CellTiter-Glo^®^ (Promega, G7572) reagent according to the manufacturer’s protocol. Briefly, cell plate was taken out of the cell culture incubator and allowed to reach room temperature for 30 minutes. After 30 minutes, a volume of CellTiter-Glo® reagent equal to the volume of cell culture medium was added to cells. Plate was covered from light, mixed for 2 minutes on an orbital shaker, and then incubated for 10 minutes in the dark. Luminescence was recorded using a PHERAstar FSX plate reader (BMG Labtech). For nuclei counts, cells were plated on 96-well PhenoPlate^TM^ microplates (PerkinElmer), treated for the stated times, and fixed in 4% paraformaldehyde (PFA, Thermo Scientific, J19943.K2) for 10 minutes. Cells were then washed, stained with DAPI (1µg/mL), and plates were imaged on a Nikon T2 microscope with automated image capture and nuclei count analysis.

### Seahorse Assays

For proliferating cells, 2,500 IMR90 cells were plated on Seahorse XFe24 cell culture microplates (Agilent, 100777-004) 24 hours prior to treatment. For oncogene-induced senescent cells, 17,500 cells were plated on a Seahorse XFe24 5 days post-transduction with HRAS G12V-encoding lentiviral particles. Media was changed every 3 days and treatment with SBI-0802162 started on day 10 post-transduction. Both proliferating and senescent IMR90 cells were treated for 3 days, at which point the Seahorse assay was performed according to the manufacturer’s protocol. BAM15 (2.5 µM) was used as the mitochondrial uncoupler, replacing FCCP.

### MitoTracker and TMRM Staining

MitoTracker Green (Cell Signaling Technologies, #9074) and TMRM (ThermoFisher Scientific, T668) were diluted to 400nM and 150nM respectively in complete DMEM media. Hoechst (AdipoGen, CDXB0030M025) was added to the same solution at a final concentration of 8.12 µM to label nuclei. Media of cells in 96-well plate was removed and replaced with media containing MitoTracker Green, TMRM, and Hoechst and plate was incubated at 37°C with 3.5% O_2_ and 5% CO_2_ for 30 minutes. After the incubation period, media was replaced with complete DMEM and cells were imaged on a Nikon T2 microscope with automated image capture.

### Bioanalytical Methods

SBI-0802162 and its stable isotope-labeled internal standard (IS), SBI-0802162-d6, were synthesized at Sanford Burnham Prebys Medical Discovery Institute (La Jolla, CA). Nicotinamide (NAM) was obtained from Sigma-Aldrich (St. Louis, MO). HPLC-grade acetonitrile (ACN), methanol, and water were purchased from Fisher Scientific (Hampton, NH). Ammonium formate (10 mM, pH 3.0) was used as the aqueous mobile phase modifier. Sample preparation: For plasma, protein precipitation was performed by adding 5 volumes of ice-cold ACN containing SBI-0802162-d6 as the IS to each plasma sample. Samples were vortex-mixed for 1 min and centrifuged at 4,000 × g for 10 min at 4°C. The supernatant was transferred to a clean 96-well plate and evaporated to dryness under a gentle nitrogen stream at 40°C. Residues were reconstituted in 50:50 (v/v) ACN:water prior to injection. For tissues, frozen tissue samples were weighed and homogenized in 4 volumes (w/v) of ice-cold 50:50 (v/v) ACN:water using a bead-based homogenizer (3 × 30 s cycles at 6,000 rpm). Homogenates were centrifuged at 14,000 × g for 10 min at 4°C. The supernatant was further processed by protein precipitation with 4 volumes of ACN containing the IS, followed by centrifugation and reconstitution as described for plasma. Final tissue concentrations were normalized to tissue wet weight (ng/g or nmol/g). Calibration standards for SBI-0802162, NAD and NAM were prepared in tissue homogenate matrix or blank mouse plasma at twelve concentration levels spanning 1–10,000 nM, using a 1/x² weighted linear regression model. The LLoQ for SBI-0802162 was 0.97 nM (S/N ≥ 10; accuracy and precision within ±20% of nominal). NAM was quantified using the same calibration curve, with IS-normalized response correcting for matrix effects and extraction variability. NAD⁺ was quantified using a separate calibration curve prepared in blank matrix; no stable isotope labeled IS was employed for NAD⁺, and concentrations are reported as absolute values normalized to the calibration curve response^84^

### LC-MS/MS Instrumentation and chromatographic conditions

Quantitative analysis was performed on an AB Sciex API 6500+ triple quadrupole mass spectrometer (AB Sciex, Framingham, MA) coupled to a Shimadzu Nexera UHPLC system (Shimadzu Corporation, Kyoto, Japan). Chromatographic separation was achieved on a Phenomenex Synergi Polar-RP column (50 × 2.0 mm, 4 µm particle size; Phenomenex, Torrance, CA) maintained at 40°C. The mobile phase consisted of (A) 10 mM ammonium formate in water (pH 3.0) and (B) ACN. A gradient elution was applied: 0–0.5 min, 5% B; 0.5–2.5 min, 5–95% B; 2.5–3.5 min, 95% B; 3.5–3.6 min, 95–5% B; 3.6–5.0 min, 5% B (re-equilibration). Total run time was 5 min at a flow rate of 0.4 mL/min, with an injection volume of 5 µL. The mass spectrometer was operated in positive ESI mode using MRM. Source parameters: ion spray voltage, 5500 V; source temperature, 500°C; curtain gas, 35 psi; GS1, 50 psi; GS2, 60 psi. MRM transitions were optimized for each analyte: SBI-0802162 and its d6-IS (compound-specific transitions), NAM (m/z 123.1 → 80.1), and NAD⁺ (m/z 664.1 → 136.1). Collision energies and declustering potentials were individually optimized by direct infusion of reference standards. Data acquisition and peak integration were performed using Analyst software v1.7 (AB Sciex).

### Mouse frailty assessments

Young (3-month-old) and aged (18-month-old) mice were exposed to control AIN-93G for 3 days prior to them being randomly assigned to each experimental group and exposed to experimental diets. Mice had ad libitum access to experimental chow and water throughout the study. Frailty assessments were performed blinded every 4 weeks as previously described^74^. Rotarod was included as part of frailty assessments. Training and assessments were performed as previously described^73^.

### Immunoblotting

For cultured cells, cells were lysed in RIPA buffer (Thermo, 89900) containing 1x protease and phosphatase inhibitors (Thermo, 78440) and 25U/mL nuclease (Thermo, 88701). For mouse tissues, samples were lysed and homogenized in the same RIPA buffer containing protease, phosphatase, and nuclease using a Bertin Precellys tissue disruptor. Lysates were then sonicated at 4°C 30 seconds on 30 seconds off for 10 cycles using the Diagenode Bioruptor Pico and were cleared by >20,000 x G centrifugation at 4°C for 10 minutes. Protein was quantified by Bradford assay (Thermo, 23246), mixed with NuPAGE LDS sample buffer (Invitrogen, NP0007), and NuPAGE reducing agent (Invitrogen, NP0009). SDS-PAGE was performed using NuPAGE 4-12% bis-tris gels (Invitrogen, NP0321BOX, NP0322BOX, WG1403BX10) run in NuPAGE MES SDS running buffer (Invitrogen, NP-0002) and transferred onto a nitrocellulose membrane using the Trans-blot turbo transfer system (Bio-Rad). Membranes were blocked with Li-Cor Intercept (TBS) blocking buffer (Li-Cor, 92760003) for 1 hour at room temperature. Primary and fluorescently labeled secondary antibodies were diluted in a 50-50 mix of Intercept (TBS) blocking buffer and PBS (Gibco, 14190-144) and imaged using the Odyssey CLx imager (Li-Cor).

### Antibodies

Antibody details are found in supplementary table 1.

### Immunofluorescence

Cells were plated on 96-well PhenoPlate^TM^ microplates (PerkinElmer), stained as described previously^85^. Following fixation, permeabilization, and blocking, cells were incubated overnight at 4°C with primary antibodies. The following antibodies were used: NAMPT (Cell Signaling Technologies, 86634S). After washing, fluorescently labeled secondary antibody was applied for 1 hour at room temperature. Cells were then stained with DAPI (1µg/mL) and plates were imaged on a Nikon T2 microscope with automated image capture.

### RNA-seq

For cultured cells, RNA was isolated using *Quick*-RNA Miniprep kit (Zymo Research, R1055) according to the manufacturer’s protocol. For mouse tissues, samples were homogenized in TRIzol (Invitrogen, 15596018) using a Bertin Precellys tissue disruptor and RNA was isolated using Direct-zol RNA miniprep (Zymo Research, R2052) according to the manufacturer’s protocol. RNA concentrations were measured using a Nanodrop One spectrophotometer (Thermo Scientific) and assessed for quality using the Agilent 4200 TapeStation system. For oncogene-induced senescent-cell studies, RNA sequencing was performed by Plasmidsaurus using Illumina Sequencing Technology with custom analysis and annotation. For irradiation-induced senescent-cell studies, library prep and sequencing was performed by the SBP genomics core. For analysis, Adapter and quality trimming was performed using Trim Galore (v0.6.7), reads were aligned to the human genome (hg38) using HISAT2 (v2.2.1), and gene-level counts were quantified using featureCounts (v2.0.3). Quality control was performed using FastQC (v0.74) and summarized with MultiQC. Differential expression analysis and normalization were conducted using DESeq2 (v2.11.40.8) with Benjamini–Hochberg correction was used to assess differentially expressed genes (genes with adjusted p-value<0.05). For mouse tissue samples, library prep and sequencing was performed by the SBP genomics core. For mouse samples, raw sequencing reads were quality-assessed using FastQC (v0.11.8) and adaptor sequences were trimmed with Trim Galore (v0.4.4). Reads were aligned to the mm10 reference genome using STAR (v2.5.3a), and gene-level counts were generated using HOMER’s analyzeRepeats.pl script. Differential expression analysis was performed using DESeq2 (v1.30.0). Genes with fewer than 10 total raw counts across all samples were excluded prior to normalization. Differentially expressed genes were defined as those with an FDR<0.05.

### Quantitative RT-PCR

RNA was isolated using *Quick*-RNA Miniprep kit (Zymo Research, R1055) according to the manufacturer’s protocol. RNA concentrations were measured using a Nanodrop One spectrophotometer (Thermo Scientific). cDNA synthesis was performed using RevertAid Reverse Transcriptase (Thermo Fischer, EP0441), Ribolock RNase inhibitor (Thermo Fischer, EO0381), and 5x RT reaction buffer (Thermo Fischer). Gene expression was quantified using PowerUp SYBR Green Master Mix (Applied Biosystems, 4485691). Primer details can be found in supplementary table 2.

### Animals

This study was approved by the Institutional Animal Care and Use Committee at Sanford Burnham Prebys MDI (AUF 23-023 and 26-013). Animals were group housed at 21-24°C, 30-70% humidity, in a 12-hour light/dark cycle under specific pathogen free conditions with ad libitum access to water and food (Teklad 2018). For details on diets utilized refer to the Animal diets in methods. For studies in figure 3 and 7E, C57BL/6J mice were purchased from Jackson Labs and switched to a base AIN-93G diet 3 days prior to study start. At study start, mice were switched to each experimental diet. For studies in figure 4 and 7A, 3-month-old C57BL/6J mice were purchased from Jackson labs and 20-month-old C57BL/6J mice were obtained from the National Institute on Aging (NIA). All mice were switched to a base AIN-93G diet 3 days prior study start. At study start mice were switched to each experimental diet for 21 days. Plasma and tissue samples were collected in the early morning and flash frozen in liquid nitrogen for analysis. For studies in 7E mice were treated with 30 or 100 mg kg^-1^ SBI-0802162 suspended in phosphate-buffered methylcellulose (0.5% methylcellulose in phosphate buffer, pH=7, 50mM) by daily oral gavage in the evening. Mice were monitored daily and weighed daily or weekly during treatment. At study endpoint, blood was collected from retro-orbital plexus and mice were euthanized by CO_2_ asphyxiation in the morning. Tissues were quickly collected and flash frozen to preserve metabolite integrity.

### Mouse diets

AIN-93G (TestDiet, 5A68) diet was supplemented with 1 g kg^-1^ chow nicotinamide (Sigma, N0636), 500 mg kg^-1^ chow SBI-0802162 (Sundia Chemicals), or a combination of both agents. All diets were generated by TestDiet.

### Blood chemistry analysis

Blood was collected from retro-orbital plexus before euthanasia in BD Microtainer K2EDTA tubes (BD, 365974) and centrifuged 2,000 x g for 10 minutes at 4°C. Plasma was collected and 100µL were loaded into VetScan Mammalian Liver Profile rotor (Zoetis). Rotor was analyzed on a VetScan VS2 Chemistry Analyzer.

### Data availability

All datasets generated in this study have been deposited in the Gene Expression Omnibus (GEO) under the following accession numbers: GSE336391, GSE336534, GSE336838.

## DECLARATION OF INTERESTS

M.A., P.D.A., M.R.J., R.R., S.J.G. are listed as inventors on a patent application filed by Sanford Burnham Prebys Medical Discovery Institute related to the combined use of NAMPT activators and nicotinamide (NAM) described in this manuscript. The remaining authors declare no competing interests.

## ACKNOWLEDGEMENTS

Work in the lab of P.D.A. was supported by P01AG092325 and U54 AG079758. M.A. was supported by the Conrad Prebys Foundation Fellowship and the Melvin and Phyllis McCardle Clause Graduate Scholarship. V.C. was supported by R35GM142495 and R21AG101323.

During the preparation of this work, the author(s) used OpenAI’s Chat GPT v5.6 to assist with proofreading and improving clarity of the manuscript. After using this tool, the author(s) reviewed and edited the content as needed and take full responsibility for the content of the published article.

## Notes

### Summary of Updates

This version corrects a typographical error in the label of Supplementary Figure 7. The correct label is Supplemental Figure 6 (Figure S6). No data, analysis, results, or conclusions have changed.

