## Supplementary figures and images for "NAMPT activation uncovers a senescence-specific vulnerability and promotes healthy aging in combination with NAM"

### Supplemental figures

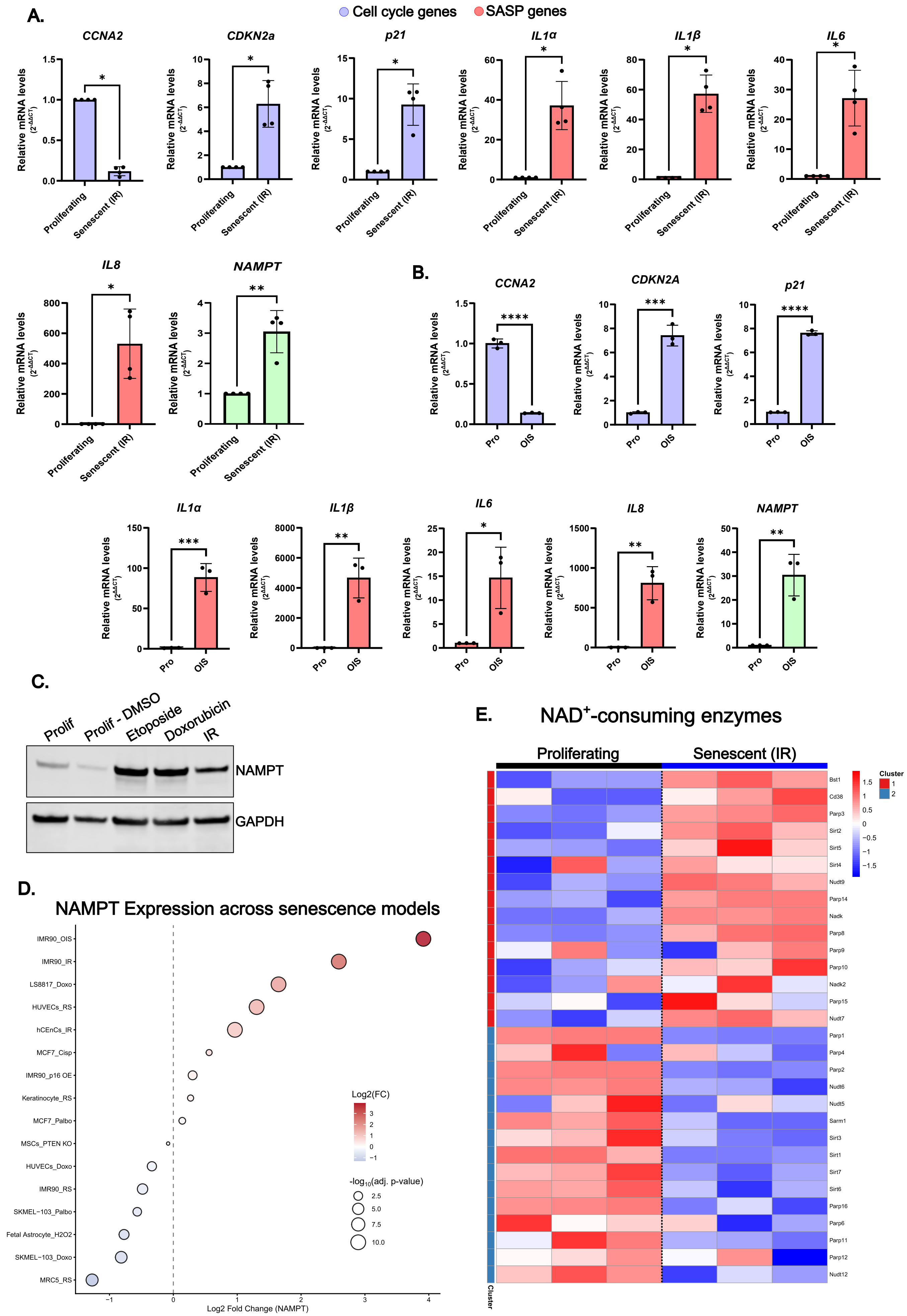

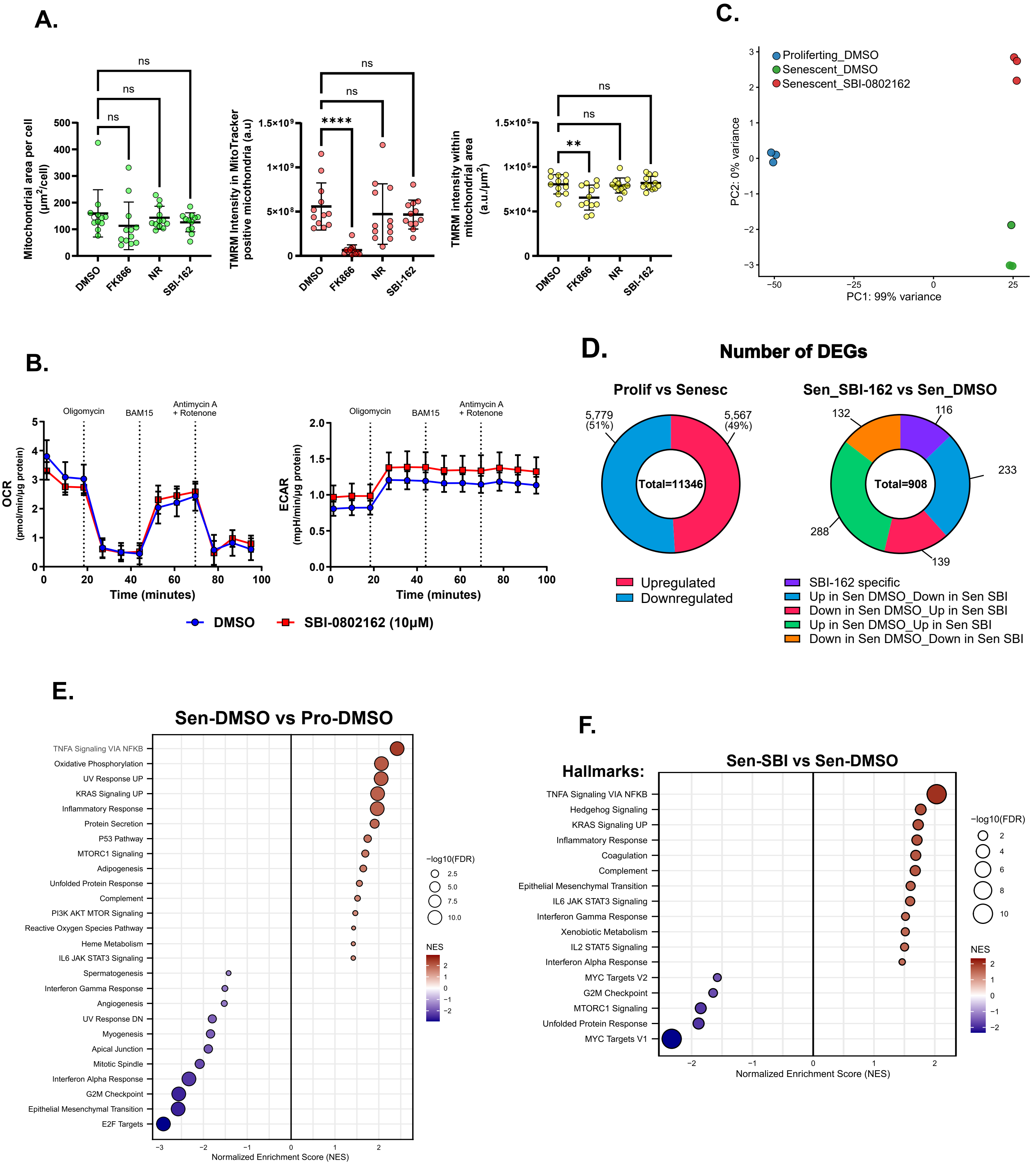

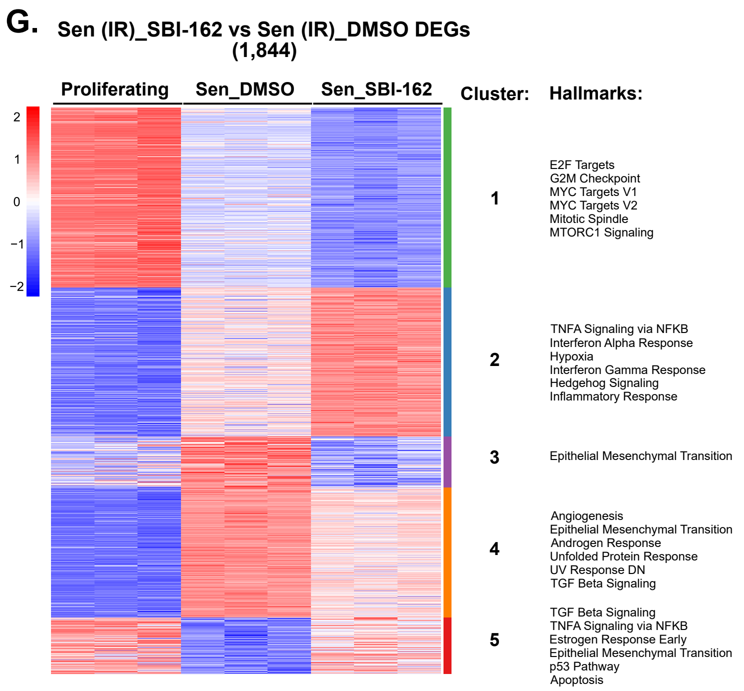

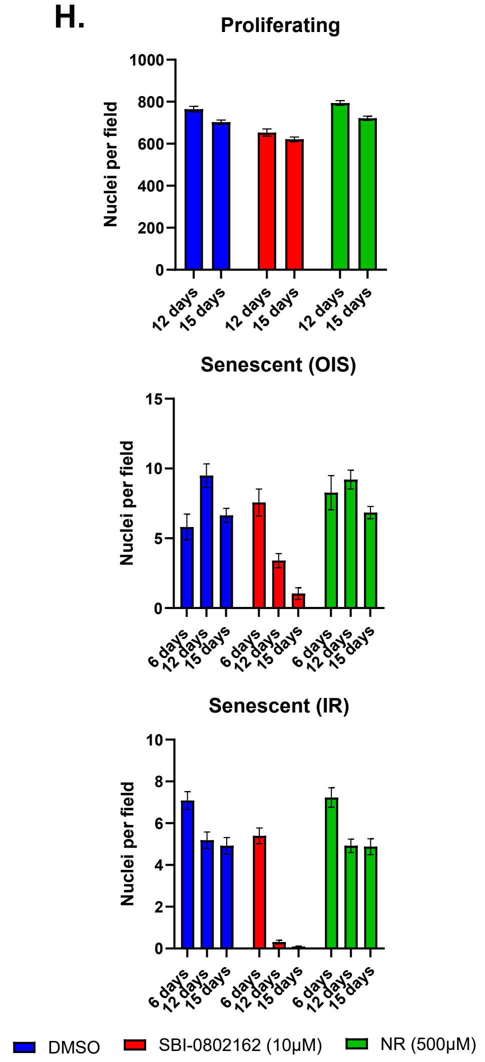

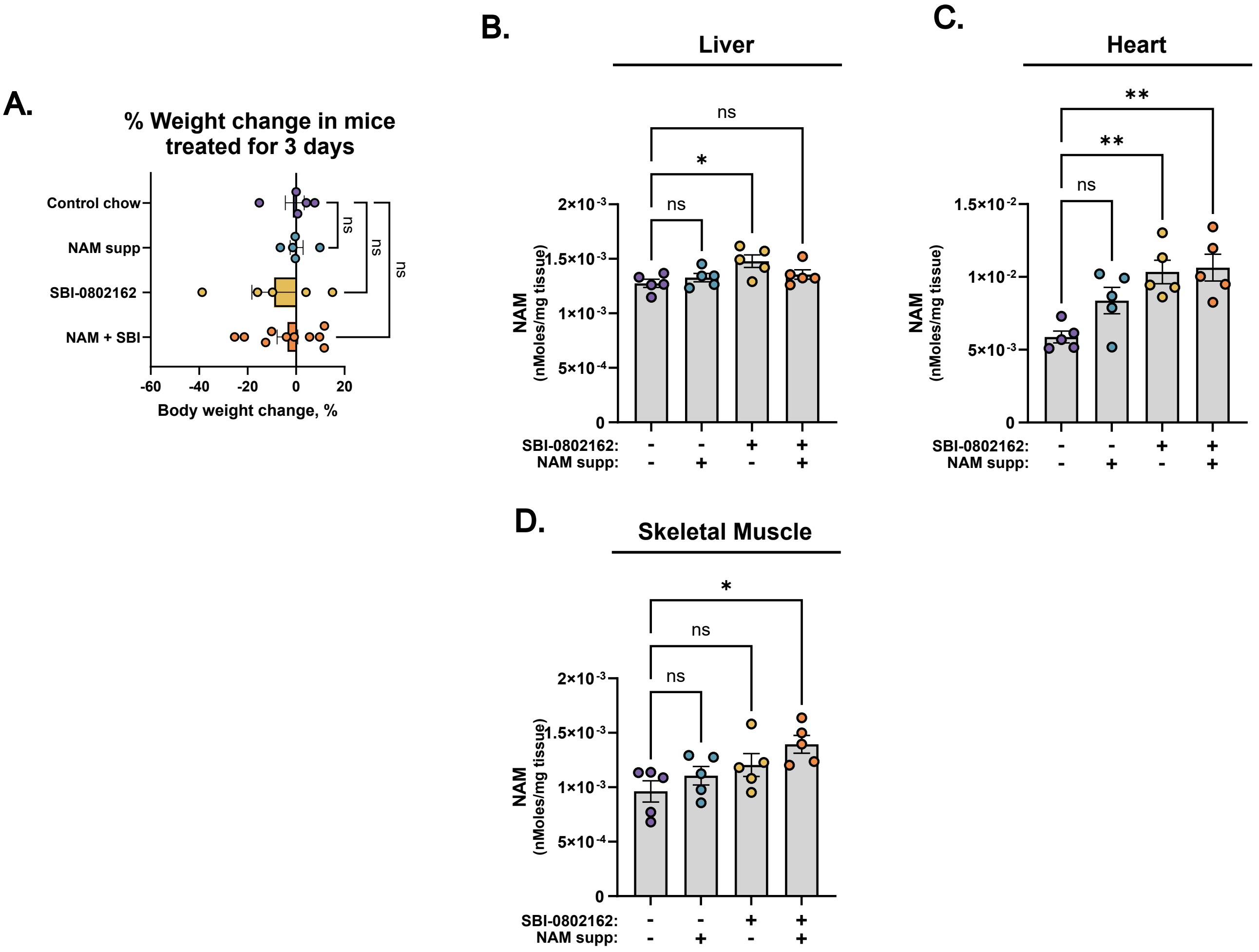

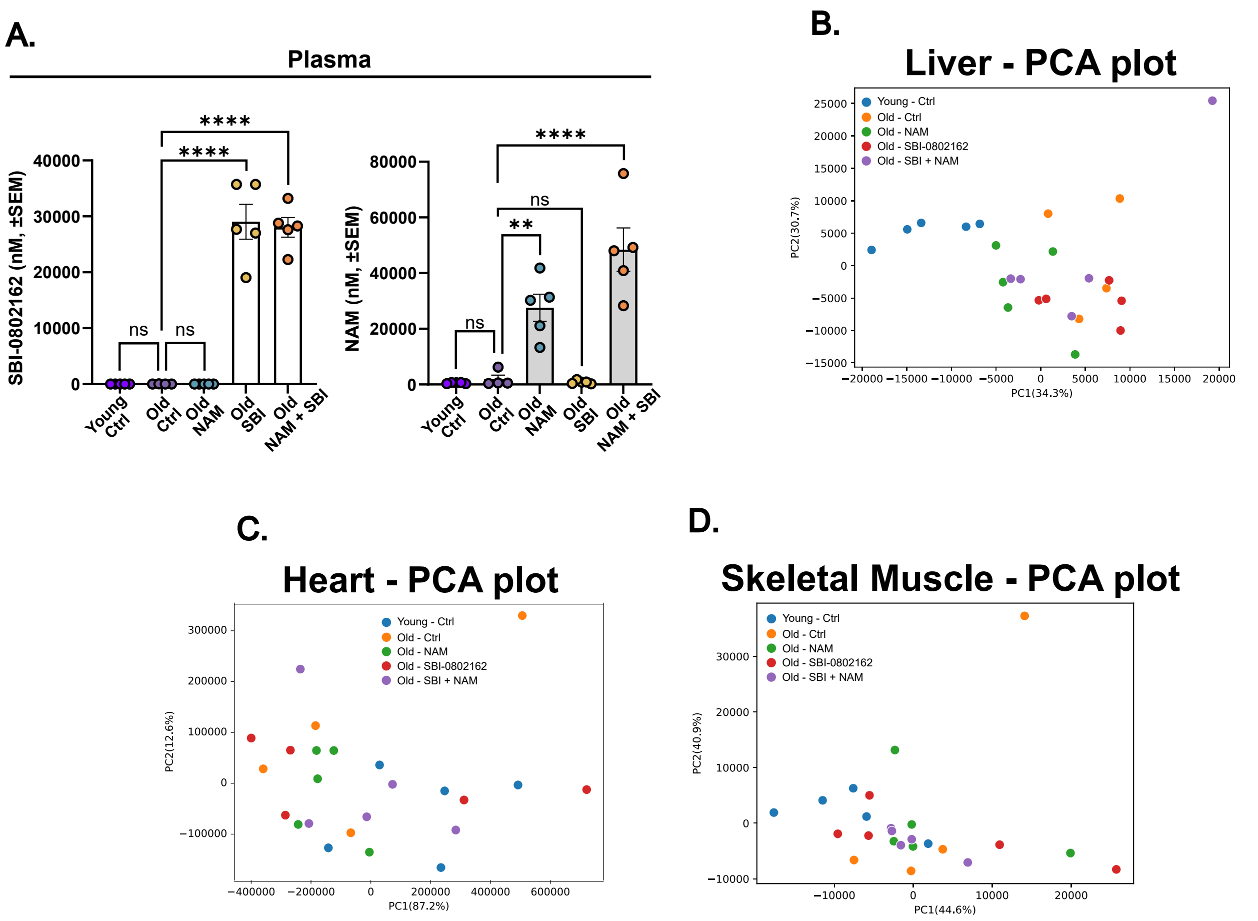

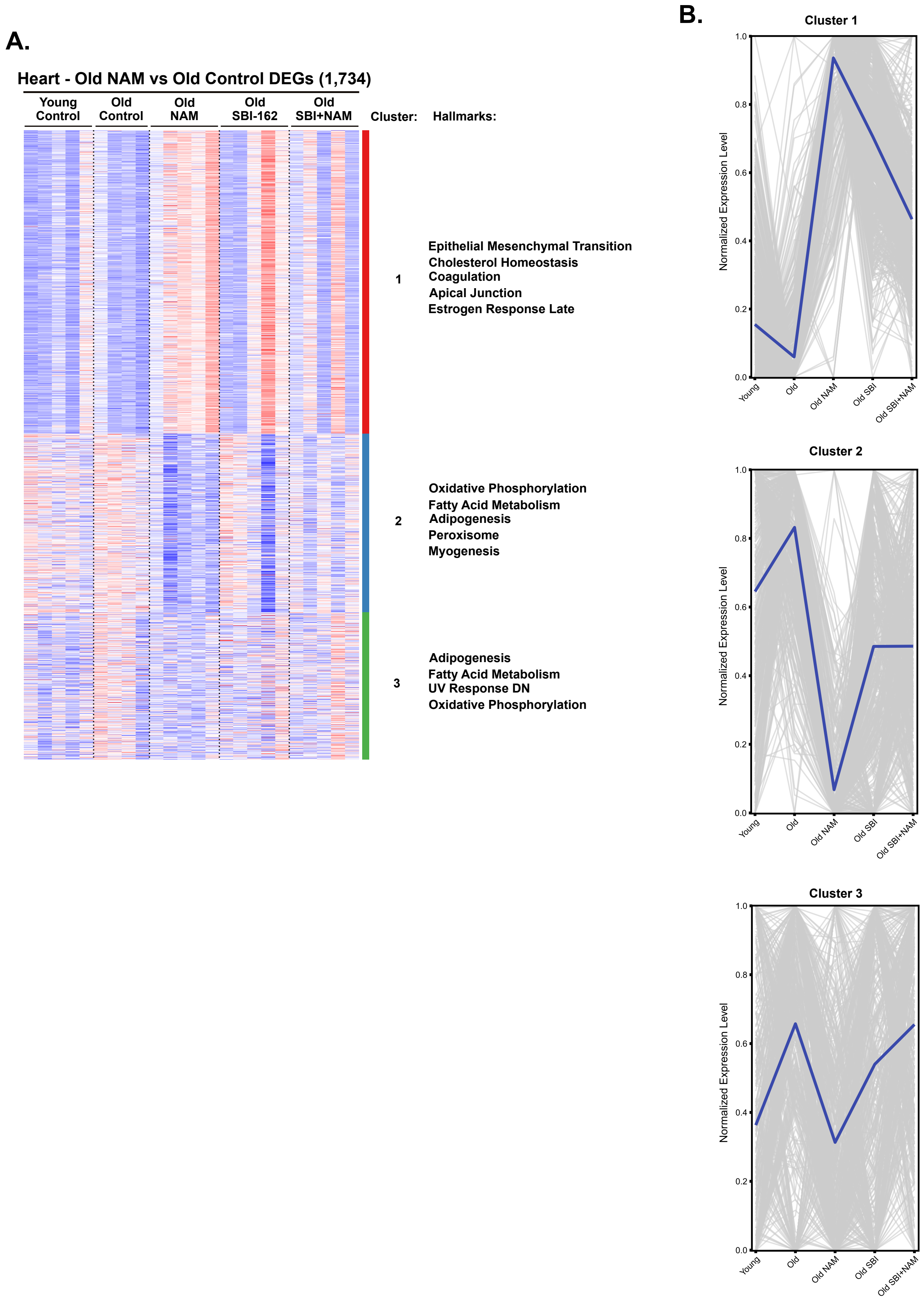

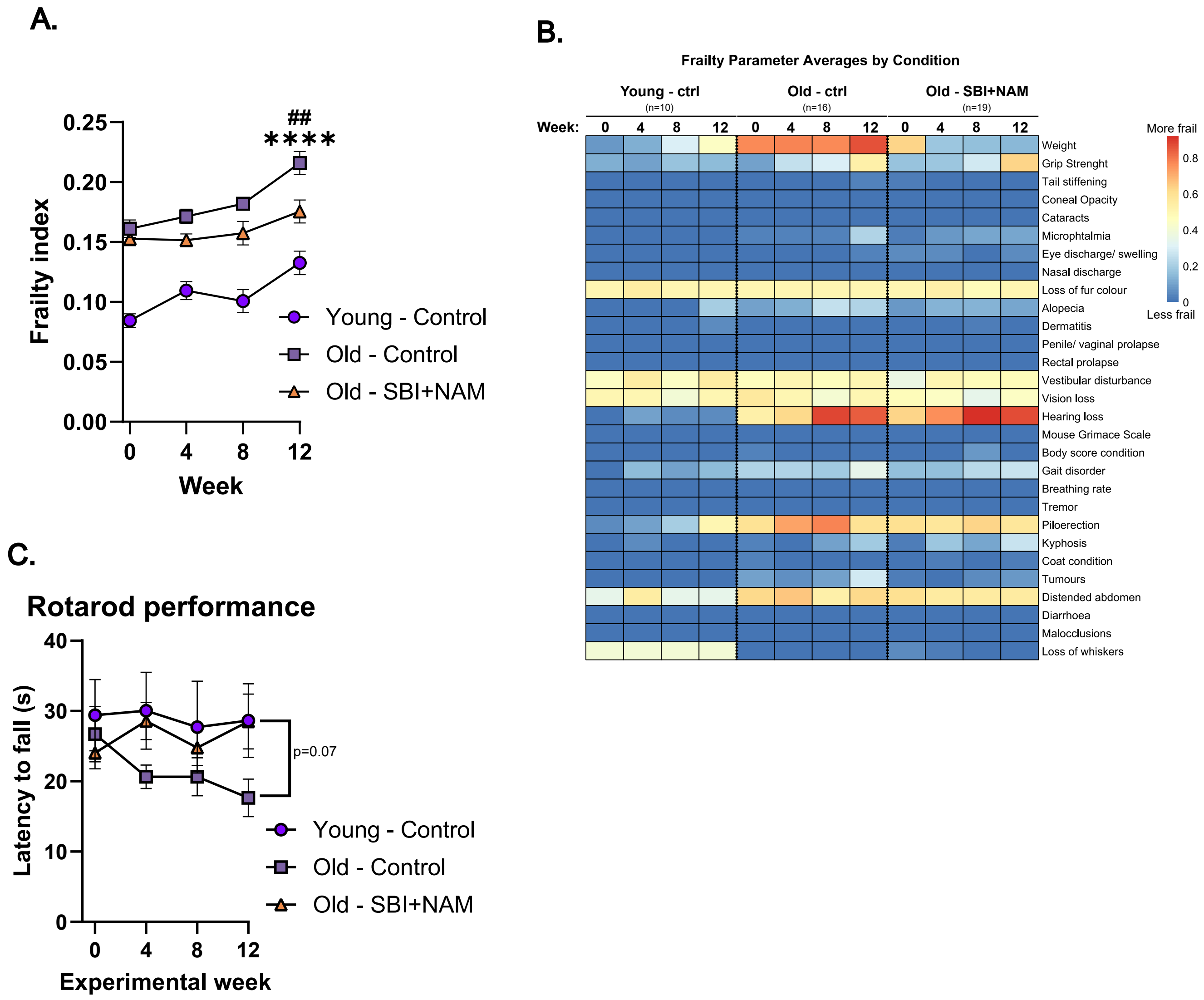
