## Supplemental figure legends for "NAMPT activation uncovers a senescence-specific vulnerability and promotes healthy aging in combination with NAM"

**Supplemental figure 1.**

**A**, Quantitative real-time (qRT) PCR of IMR90 cells 15 days post-irradiation. **B**, qRT-PCR of OIS cells to confirm senescent phenotype. **C**, NAMPT protein levels in proliferating, proliferating cells treated with DMSO, etoposide-induced, doxorubicin-induced, and IR-induced senescent cells. **D**, Dot plot showing NAMPT expression across independent publicly available RNA-seq datasets from multiple cell types and senescence-inducing stimuli. **E**, Expression level of 30 major $NAD^+$ consuming enzymes in proliferating and irradiation-induced senescent cells. Each dot in S1A and S1B represents a biological replicate. Error bars denote mean ± s.d. Statistical analysis was performed using unpaired parametric T-tests. ns = not significant, *p<0.05, **p<0.01, ***p<0.001, ****p<0.0001.

**Supplemental figure 2.**

**A**, Mitochondrial area and membrane potential analysis in proliferating IMR90 after treatment with FK866 (0.1 µM), NR (500 µM), or SBI-0802162 (10 µM) for 96 hours. **B**, Seahorse OCR (left) and ECAR (right) traces in proliferating IMR90 cells treated with DMSO or SBI-0802162 (10 µM) for 96 hours. OCR and ECAR were normalized to total protein in each replicate. **C**, Principal component analysis of RNA-sequencing profiles from proliferating, DMSO-treated senescent cells, and oncogene-induced senescent cells treated with SBI-0802162 (10 µM) for 96 hours. **D**, Donut plots showing the number and direction of differentially expressed genes between proliferating and senescent cells (left), and between SBI-0802162 treated and DMSO treated oncogene-induced senescent cells (right). **E**, GSEA pathway enrichment analysis of genes differentially expressed between oncogene-induced senescent and proliferating cells. **F**, GSEA pathway enrichment analysis of genes differentially expressed between SBI-0802162-treated and DMSO-treated oncogene-induced senescent cells. **G**, Heatmap of differentially expressed genes between SBI-0802162 and DMSO-treated irradiation-induced senescent cells, shown alongside proliferating cells for comparison. Major enriched Hallmark pathways within each gene cluster are indicated to the right of the heatmap. **H**, Quantification of nuclei per field in proliferating, irradiation-induced, and oncogene-induced senescent cells treated every 3 days with DMSO, SBI-0802162 (10 µM), or NR (500 µM). Each dot in plots represents a biological replicate, except in S2A where each dot represents a single frame from 3 biological replicates were analyzed. Error bars denote mean ± s.d. Statistical analysis was performed using One-Way ANOVA and Dunnett's multiple comparisons test. ns = not significant, *p<0.05, **p<0.01, ***p<0.001, ****p<0.0001.

**Supplemental figure 3.**

**A**, Percent body weight change in mice after 3 days of dietary treatment with control chow or diets containing NAM, SBI-0802162, or combined SBI-0802162 and NAM. B, NAM levels in liver, heart (**C**), and skeletal muscle in mice exposed to the experimental diets in figure 3D. Each dot in plots represents an individual mouse. Error bars represent mean ±

SEM. Statistical analysis was performed using One-Way ANOVA and Dunnett's multiple comparisons test. ns = not significant, *p<0.05, **p<0.01, ***p<0.001, ****p<0.0001.

**Supplemental figure 4**

A, Plasma levels of SBI-0802162 and NAM after 21 days of dietary treatment. **B**, Principal component analysis of RNA-sequencing profiles from liver, heart (**C**), and skeletal muscle (**D**) after in-chow dosing with NAM, SBI-0802162, or combined SBI-0802162 plus NAM for 21 days. Each dot in plots represents an individual mouse. Error bars represent mean ± SEM. Statistical analysis was performed using One-Way ANOVA and Dunnett's multiple comparisons test. ns = not significant, *p<0.05, **p<0.01, ***p<0.001, ****p<0.0001.

**Supplemental figure 5.**

**A**, Heatmap of differentially expressed genes in the heart between NAM-supplemented and age-matched control mice shown alongside all experimental conditions depicted in diagram in Figure 4A. Major enriched hallmark pathways within each gene cluster are indicated to the right of the heatmap. Each column in heatmap represents an individual mouse. **B**, Normalized average gene expression level within each cluster in Figure S5A.

**Supplemental figure 6.**

**A**, Average frailty index scores measured longitudinally over 12 weeks in young control, aged control and aged mice treated with combined SBI-0802162 plus NAM. Higher values indicate increased frailty. **B**, Heatmap showing average scores for individual frailty parameters across young control, aged control, and aged combination-treated mice at baseline and weeks 4, 8, and 12. Higher values indicate increased frailty. **C**, Rotarod performance measured as latency to fall over the 12-week study. Data shown as mean ± SEM. Statistical analysis was performed using One-Way ANOVA and Dunnett's multiple comparisons test. For S7A # indicate comparison between old-control and old-SBI+NAM and * represent comparison between young control and old control. ns = not significant, *p<0.05, $^{**/\#\#}$p<0.01, ***p<0.001, ****p<0.0001.
