## Supplemental table 1 for "NAMPT activation uncovers a senescence-specific vulnerability and promotes healthy aging in combination with NAM"

**Supplemetal Table 1**

| Antibody target | Host | Clonality | Catalog number | Vendor | Concentration for western blot | Concentration for IF |
|---|---|---|---|---|---|---|
| NAMPT | Rabbit | mAb | 86634S | Cell Signaling Technologies | 1:1000 | 1:200 |
| phospho Rb (S807/811) | Rabbit | mAb | 8516S | Cell Signaling Technologies | 1:1000 | N/A |
| p16 | Rabbit | pAb | 10883-1-AP | Proteintech | 1:1000 | N/A |
| GAPDH | Mouse | mAb | SC-47724 | Santa Cruz | 1:5000 | N/A |
| p16 | Rabbit | mAb | Ab211542 | Abcam | 1:1000 | N/A |
| Vinculin | Mouse | mAb | 66301-1-LG | Proteintech | 1:1000 | N/A |
| PARP1 | Rabbit | pAb | 9542S | Cell Signaling Technologies | 1:1000 | N/A |
| SIRT1 | Mouse | mAb | 8469S | Cell Signaling Technologies | 1:1000 | N/A |
| SIRT2 | Rabbit | pAb | 19655-1-AP | Proteintech | 1:1000 | N/A |
| SIRT3 | Rabbit | mAb | 5490S | Cell Signaling Technologies | 1:1000 | N/A |
| STAT1 | Rabbit | pAb | 9172S | Cell Signaling Technologies | 1:1000 | N/A |
| phospho STAT1 (Y701) | Rabbit | mAb | 9167S | Cell Signaling Technologies | 1:1000 | N/A |
| Lamin B1 | Rabbit | pAb | 12987-1-AP | Proteintech | 1:1000 | N/A |
| CCND2 | Rabbit | Recombinant | 3741 | Cell Signaling Technologies | 1:1000 | N/A |
| CCND1 | Rabbit | Recombinant | MA5-16356 | Invitrogen Antibodies | 1:1000 | N/A |
| CCNB1 | Rabbit | mAb | 04-220 | Millipore Sigma | 1:1000 | N/A |
| IRDYE® 680RD Anti-mouse | Donkey | pAb | 926-68072 | LICOR bio | 1:15000 | N/A |
| IRDYE® 800CW Anti-rabbit | Donkey | pAb | 926-32213 | LICOR bio | 1:15000 | N/A |
| Goat anti-Rabbit IgG (H+L) | Goat | pAb | A11008 | Fisher Scientific | N/A | 1:500 |
