## Supplemental table 2 for "NAMPT activation uncovers a senescence-specific vulnerability and promotes healthy aging in combination with NAM"

| Primer name | Target | Sequence (5'→3') |
|---|---|---|
| GAPDH_F | Human GAPDH | CATGTTCGTCATGGGTGTGAACCA |
| GAPDH_R | Human GAPDH | AGTGATGGCATGGACTGTGGTCAT |
| RPL13_F | Human RPL13 | GTCTGAAGCCTACAAGAAAGTTTGC |
| RPL13_R | Human RPL13 | CGTTCTTCTCGGCCTGTTTCC |
| CCNA2_F | Human CCNA2 | CTCTACACAGTCACGGGACAAAG |
| CCNA2_R | Human CCNA2 | CTGTGGTGCTTTGAGGTAGGTC |
| p21_F | Human p21 | GGATGTCCGTCAGAACCCAT |
| p21_R | Human p21 | GTGGGAAGGTAGAGCTTGGG |
| CDKN2A_F | Human CDKN2A | CTGCCCAACGCACCGAATAG |
| CDKN2A_R | Human CDKN2A | CCACCAGCGTGTCCAGGAAG |
| IL1α_F | Human IL1α | AGTGCTGCTGAAGGAGATGCCTGA |
| IL1α_R | Human IL1α | CCCCTGCCAAGCACACCCAGTA |
| IL1β_F | Human IL1β | TGCACGCTCCGGGACTCACA |
| IL1β_R | Human IL1β | CATGGAGAACACCACTTGTTGCTCC |
| IL8_F | Human IL8 | GAGTGGACCACACTGCGCCA |
| IL8_R | Human IL8 | TCCACAACCCTCTGCACCCAGT |
| IL6_F | Human IL6 | AAATTCGGTACATCCTCGACG |
| IL6_R | Human IL6 | TTTCACCAGGCAAGTCTCC |
| NAMPT_F | Human NAMPT | CGGCAGAAGCCGAGTTCAA |
| NAMPT_R | Human NAMPT | GCTTGTGTTGGGTGGATATTGTT |
